# Chromosome X-wide analysis of positive selection in human populations: from common and private signals to selection impact on inactivated genes and enhancers-like signatures

**DOI:** 10.1101/2021.05.24.445399

**Authors:** Pablo Villegas-Mirón, Sandra Acosta, Jessica Nye, Jaume Bertranpetit, Hafid Laayouni

## Abstract

The ability of detecting adaptive (positive) selection in the genome has opened the possibility of understanding the genetic bases of population-specific adaptations genome-wide. Here we present the analysis of recent selective sweeps specifically in the X chromosome in different human populations from the third phase of the 1000 Genomes Project using three different haplotype-based statistics. We describe numerous instances of genes under recent positive selection that fit the regimes of hard and soft sweeps, showing a higher amount of detectable sweeps in sub-Saharan Africans than in non-Africans (Europe and East Asia). A global enrichment is seen in neural-related processes while numerous genes related to fertility appear among the top candidates, reflecting the importance of reproduction in human evolution. Commonalities with previously reported genes under positive selection are found, while particularly strong new signals are reported in specific populations or shared across different continental groups. We report an enrichment of signals in genes that escape X chromosome inactivation, which may contribute to the differentiation between sexes. We also provide evidence of a widespread presence of soft-sweep-like signatures across the chromosome and a global enrichment of highly scoring regions that overlap potential regulatory elements. Among these, enhancers-like signatures seem to present putative signals of positive selection that might be in concordance with selection in their target genes. Also, particularly strong signals appear in regulatory regions that show differential activities, which might point to population-specific regulatory adaptations.

## INTRODUCTION

The evolution of *Homo sapiens* has been strongly shaped by positive selection in the last 100,000 years, by adaptations to specific environments, diets, and cognitive challenges that modern human populations encountered as they expanded across the globe. Surviving such challenges has left remarkable footprints of selection in the human genome, like in the lactase (*LCT*) locus in European populations (Bersaglieri et al., 2004; Wang et al., 2020), genes involved in skin pigmentation like *MC1R* (John et al., 2003) or genes implicated in resistance to severe malaria infection like *CD40L* and *G6PD* (Sabeti et al., 2002). Studying the evolutionary processes that resulted from these adaptations can uncover which path our ancestors travelled along to give rise to extant adaptations of present human populations.

The development of new methods to study recent selection in natural populations (Fan et al., 2016; Field et al., 2016; Pavlidis et al., 2017) has settled genomic selection scans as one of the main approaches to study the genetic origin behind such adaptations (Mathieson et al., 2015; Casillas et al., 2018; Lopez et al., 2019; Walsh et al., 2020). However, most of these scans have focused on coding regions as the main target of selection and have attached greater importance to the study of processes driven by *de novo* mutations, that leave strong and more evident selection signatures (classical hard sweeps). Although gene regulation is considered to be the primary driver of phenotypic changes in the evolution of *Homo sapiens* (King and Wilson, 1975), these strategies might have overlooked standing variation in regulatory regions as the main targets of rapid adaptations, which seem to be more likely selection targets and are marked by more subtle signatures, like soft sweeps (Fu and Akey, 2013; Scheinfeldt and Tishkoff, 2013; Messer and Petrov, 2013).

Selection on standing variation seems to be a more common mode of selection and soft sweeps a more widespread signature in human genomes (Hernandez et al., 2011; Schrider and Kern, 2017). Multiple modes of selection can originate a soft sweep signature: on standing variation, de novo mutation on multiple haplotypes and recurrent origination of adaptive alleles (Schrider et al., 2015; Hermisson and Pennings, 2017). However, sometimes patterns of variation might exhibit different degrees of “softness” and, together with confounding factors like demography or recombination, display sweep-like signatures where the picture is not clear enough so as to define a region under a specific selection regime (Messer and Petrov, 2013). Therefore, sometimes it is difficult to differentiate signatures due to hard or soft sweeps, and often linked regions under selection may present properties of both types of signals (Schrider et al., 2015).

The X chromosome, although it’s been studied in terms of recent positive selection in human populations (Casto et al., 2010; Veeramah et al., 2014; Johnson and Voight, 2018), remains to be addressed more completely, including selection on regulatory regions and a focused analysis of signatures of selection on standing variation. The X and Y chromosomes have different inheritance models than the autosomes as well as different effective population sizes, making the outcome of selection pressures inconsistent to the rest of the genome. In order to study the X chromosome, these different properties have to be taken into account and, to analyse the selection events that took place, chromosome-specific demographic models and region-specific recombination maps must be incorporated to approximate a more realistic scenario.

The unique properties of the X chromosome, as compared with autosomes, have been extensively studied (Vicoso and Charlesworth, 2006; Mank et al., 2016; Meisel and Connallon, 2013). Dosage compensation of the X chromosome, the process that allows XY males and XX females to cope with different gene copy numbers on the X, might lead to sex-specific patterns of selection. This process involves the random transcriptional silencing of one of the X chromosomes in females. However, this inactivation is not complete for all the genes. Evidence suggests that around 23% of the X-linked genes “escape” inactivation and express both chromosomal copies (Balaton et al., 2015; Tukiainen et al., 2017), leading to a sex-biased expression of these genes, which might be responsible for dimorphic traits, and potentially, adaptations associated with phenotypic diversity. Despite the few studies of selection on these genes, some evidence indicates that these regions have been under purifying selection (Park et al., 2010). Thus, it is of interest to see whether positive selection has operated in these regions and the relative importance that inactivation may have on the process of natural selection.

The faster-X hypothesis (Meisel and Connallon, 2013) postulates that selection occurs faster in genes on the X than in autosomes due to the hemizygosity of males, this is supported by recent evidence that found increased selection levels in the sexual chromosome (Veeramah et al., 2014). Moreover, different effects of mutations in males versus females have been well-established (Vicoso and Charlesworth, 2006). The difference in the replication rate between female and male germ lines favours this hypothesis. The higher probability of suffering consequences due to deleterious and adaptive mutations most likely has led to a different selection process. Altogether, these factors may lead to specific patterns which reflect the sex-biased evolution in humans.

In this study, we conduct a selection scan on the X chromosome of 15 human populations from three different continental groups (Sub-Saharan Africa, Europe and Asia). We sought to identify signatures of recent positive selection by considering hard and soft sweeps. potentially affecting both coding and non-coding regions. With this we aim to disentangle how positive selection has shaped the diversity patterns in the X chromosome across the globe.

## MATERIALS AND METHODS

### Genetic Data

Phased VCF files from the third phase of the 1000 Genomes Project were downloaded from the project database (Auton et al., 2015). These data are whole-genome (mean depth of 7.4X) and targeted exome sequences (mean depth of 65.7X) with a total of 2,504 individuals across 26 different populations, covering three continental groups. Due to the methodological complexity, only the non-admixed populations of each geographical group were analysed. In Africa: Esan (Nigeria, ESN), Gambian (Wester Divisions in the Gambia, GWD), Luhya (Webuye, Kenya, LWK), Mende (Sierra Leone, MSL), Yoruba (Ibadan, Nigeria, YRI); Europe: Utah residents with northern and western European ancestry (CEU), Finnish (Finland, FIN), British (England and Scotland, GBR), Iberians (Spain, IBS), Toscani (Italy, TSI); and Asia: Chinese Dai, (Xishuangbanna, China, CDX), Han Chinese (Beijing, China, CHB), Southern Han Chinese (China, CHS), Japanese (Tokyo, Japan, JPT), Kinh (Ho Chi Minh City, Vietnam, KHV). We applied filters to remove duplicated variants found in the X chromosome. These errors were reported to the 1000 Genomes Project (www.1000genomes.org).

The X chromosome consists of both pseudoautosomal regions (PAR) and non-pseudoautosomal regions (nPAR). Since the PAR behaves differently and does not follow the same inheritance rules than the rest of the X chromosome, we removed these regions keeping only bi-allelic variants within the position range of the nPAR region (~2.7-155.0 Mb) (Flaquer et al., 2008).

We reformatted the VCF file so that the ancestral allele was the reference and the derived allele was the alternative. The human ancestral alleles determined by their state in chimpanzee were downloaded from the 1000 Genomes Project mapped to human reference GRCh37. We removed any SNP whose ancestral status was unknown, resulting in a total of 2.852.479 SNPs from 1511 individuals (504 Africans, 503 Europeans, and 504 Asians).

We downloaded a population-combined genetic map of the nPAR region (http://mathgen.stats.ox.ac.uk). This map was based on the first phase of The 1000 Genomes Project (GRCh37). In order to use the map for phase three data, we performed a linear interpolation of the missing values using the command *approx* from the statistical programming language R (R Core Team, 2020).

### Neutral simulations

We used the *msms* software (Ewing and Hermisson, 2010) to simulate neutral scenarios. For the X chromosome we implemented a three-population demographic neutral model adapted from Henn et al. (2015) for the continental populations Africa (AFR), Europe (EUR), and Asia (ASI) with a mutation rate of 1.25×10^-8^ mutations per base per generation (Henn et al., 2015), a generation time of 30 years, a recombination rate of 1.3×10^-8^ per nucleotide, and a Watterson estimator θ (4Neμ) of 328.79. We chose a three-population model due to the high similarity within continents, with a mean sample size of AFR: 152, EUR: 153, and ASI: 149. Since the effective population size of the X is ¾ the size of the autosomes, we accounted for this by modifying the population sizes, resulting in N_e_ for AFR: 23220, EUR: 2479, and ASI: 907. We simulated multiple regions of 600 kb in order to reproduce the total length of the X chromosome, by using the following parameters:

~~~
msms -N 10538.25 -ms 454 254 -t 316.1475 -r 328.7934 600000 -I 3 152 153 149
0 -n 1 2.204 -n 2 3.2542 -n 3 7.4055 -g 2 56.61 -g 3 96 -ma x 0.3542 0.1462
0.3542 x 1.3562 0.1462 1.3562 x -ej 0.0464 3 2 -en 0.0464 2 0.2939 -em 0.0464
1 2 4.9314 -em 0.0464 2 1 4.9314 -ej 0.14022 2 1 -en 0.364 1 1 -oTPi 30000
25000 -tt -oAFS
~~~

In order to contrast the results obtained for the X chromosome, we analysed the complete set of autosomes in the human genome. The same procedure to detect positive selection as for the X was followed. To do so we performed the appropriate autosomal neutral simulations and used the percentile 99^th^ as extreme distribution cut-off to compare the regions under positive selection. Also, the Refseq gene annotation from the UCSC database table browser (Karolchik et al., 2004) (downloaded June 2020) was considered.

### Scan for signals of selection

Advances in the statistics used to detect selective sweeps allow for the analysis of linkage disequilibrium decay (Pybus et al., 2015, Biswas and Akey, 2006; Vallender, 2004; Sabeti et al., 2006; Garud et al., 2015). These methods rely on detecting decreased variation surrounded by a region with high linkage disequilibrium (LD). The LD increases and the variation decreases as the frequency of the selected allele rises in the population. Once the selected allele is fixed, selection will relax, allowing for variation to recover through new mutations and recombination. The extended haplotype homozygosity (EHH) computes the probability that, at a given distance from a core region, two randomly chosen chromosomes carry homozygous SNPs for the entire interval. In this analysis we made use of three different haplotype-based statistics that rely on the EHH computation at a tested SNP, taking into account the ancestral and derived allele state.

The integrated haplotype score (iHS) is the integral (Voight et al., 2006) of EHH and is designed to detect incomplete hard sweeps. These are signatures of recent, ongoing selection that are characterized for presenting long blocks of homozygosity found in haplotypes with a high frequency of derived alleles. We have used two methods to detect signatures that resemble soft sweeps. The integrated haplotype homozygosity pooled (iHH12) (Torres et al., 2018) is an adaptation of the H12 statistic by Pickrell et al. (2018) and is able to detect signatures of both hard and soft sweeps, and the number of segregating sites by length (nSL) (Ferrer-Admetlla et al., 2014), a modification of iHS with a higher robustness to recombination rate variation and with an increased power to detect soft sweeps. These are the footprints left by selection processes that target variants at intermediate frequencies. On the contrary to the hard sweeps, that involve the fixation of a single *de novo* mutation due to a specific environmental change, soft sweeps might be generated by the selection of an allele that was drifting neutrally at the moment of the change. Also, these footprints might appear when different alleles are selected simultaneously at the same locus. Therefore, the footprints left by this kind of process are not as evident as the signatures left by the hard sweeps, since the diversity reduction left by the sweep is lower. These tests for recent positive selection are standardized (mean 0, variance 1) by the distribution of observed scores over a range of SNPs with similar derived allele frequencies. Here, we use the three tests, iHS, nSL and iHH12, to detect selective sweeps in the X chromosome.

The candidate signals for selection may point to putative targets of recent selection which are of particular interest in the study of human evolution and may help to understand complex phenotypes of medical relevance. The calculations of iHS, nSL and iHH12 were computed with the software *selscan* (Szpiech and Hernandez, 2014), an application that implements different haplotype-based statistics in a multithreaded framework. We allow for a maximum gap of 20kb and keep only SNPs with a minor allele frequency (MAF) higher than 5%. These parameters reduce the number of false positives due to the presence of gaps in the data, however special care must be taken when interpreting these results since false positive rate could increase with other confounding factors. The same procedure was applied on the simulated data in order to compare the empirical distributions with a neutral score background. The standardization was performed by the *norm* function within the *selscan* package for each population and test separately. The calculations of the Tajima’s D scores in Figure 2 were performed by using the software package *VCFtools* (0.1.14) with a non-overlapping 10kb sized window-based approach (Danecek et al., 2011).

The program *selscan* considers ancestral and derived alleles separately. iHS and nSL report positive values when the derived allele is selected while negative values indicate the ancestral allele is favoured (Szpiech and Hernandez, 2014). iHH12 makes no distinction between the two allele states. Since a sweep may also be produced by the hitchhiking of ancestral alleles with the selected variant, absolute values were considered. The per-SNP scores were summarized by using a position-based sliding window approach of size 20kb with a 20% overlap (4kb). Windows with 20 SNPs or fewer were removed. The mean scores were calculated in each test in order to interpret the presence or absence of a selective sweep. To search for candidate windows under positive selection, we compared the distributions of the summary observed values to the simulations and considered 99^th^ and 99.9^th^ percentiles in the simulated distribution as critical values to have evidence against neutrality. Empirical summary values over these thresholds were considered as putative signals of positive selection. No p-values were associated with the significance of these windows.

The haplotype structure in regions under putative positive selection was determined with the program *fastPHASE* (Scheet and Stephens, 2006). This software applies a Hidden Markov Model (HMM) on haplotype data to obtain the frequencies of a certain SNP to be in a haplotype cluster according to the similarity between them, such that the region is divided into a mosaic of clusters per population that reflects the patterns of haplotypic variation.

In order to assess commonalities and differences across populations, we identified the regions under selection that are in the extreme tail of more than one population. Since a region under positive selection can be captured by more than one test depending on the variable degree of “softness” in its locus, the shared sweeping regions were constructed by using the candidate windows reported in the extreme 99^th^ percentile across the three selection tests. Sweeping regions that overlap across more than two populations of the same continental group were considered shared in that group.

### Gene Ontology

We downloaded the Refseq gene annotations from the UCSC database table browser (Karolchik et al., 2004) in June 2020 to annotate the X chromosome. This annotation describes all the transcripts including 5’ and 3’ untranslated regions (UTR), coding, and non-coding genes. We merged these annotations with our empirical data using *Bedtools intersect* (Quinlan and Hall, 2010). We intersected our candidate windows under selection with the annotated genomic regions to obtain a list of genes under putative positive selection. Finally, an Overrepresentation Enrichment Analysis (OEA) was performed on the most extreme top 100 genes for each population with the online tool *WebGestalt GSAT* (Gene Set Analysis Toolkit). The multiple testing was adjusted using the Benjamini-Hochberg correction, accepting ontology terms with a global false discovery rate (FDR) ≤ 0.05 as significant.

In order to focus on putative regions with the highest selection scores, we selected the top windows that fall into the 99.9^th^ percentile. The SNPs contained in these windows were annotated using the *ANNOVAR* program (Wang et al., 2010), which aggregates the UCSC annotations: GWAS Catalog, CADD scores, GERP++ scores, Conserved transcription factor binding sites (TFBS) in the human/mouse/rat alignment, segmental duplications, and clusters of TFBS based on ChIp-seq data. In order to identify the most interesting SNPs inside each region, we considered SNPs with an individual selection value within the 1% extreme tail of the distribution (|iHS| and |nSL| ~ 2.5 in all populations, and iHH12 ~ (Africa: 4.1, Europe: 3.8 and Asia: 3.6)) and a PHRED-scaled CADD score ≥ 10, which represents the whole genome 1% most deleterious SNPs according to Kircher et al. (2014). Also, as a way to prioritize SNPs located in regulatory regions, we explored the potential effects of SNPs from both 99th and 99.9th top windows within functional regions by using RegulomeDB (Boyle et al., 2012). This database uses ENCODE data sets to annotate variants that are likely to belong to a functional region and thus suggest possible hypotheses to the SNPs within the selection signal. This database presents a classification scheme that scores the variants according to the support they have of functional elements. The functional categories decrease with the relevance of each variant. In this line, the category 1 corresponds with those variants that present an eQTL and support from other ENCODE data, while the category 6 only presents a hit in a single motif.

### “Escape” genes selection analysis

The putative selection signals were used to explore potential signatures of recent positive selection in genes with X chromosome inactivation (XCI) status. Several studies have cataloged XCI gene status in order to categorize genes that escape inactivation (Balaton et al., 2015; Carrel and Willard, 2005; Cotton et al., 2013). For this analysis, the inactivation status was considered using the catalog by Tukiainen et al. (2017), which includes a consensus of XCI statuses from previous studies (Carrel and Willard, 2005; Cotton et al., 2013) and extends it by creating a landscape of human XCI across different tissues (GTEx project, v6p release) and individuals. The integrated statuses of these studies fall into three categories: *escape* (if “escape” and “variable”), *variable* (if “escape” and “inactive”), and *inactive* (if “variable” and “inactive”). Contingency tables were constructed based on selection (Selected/Not selected) and XCI (Escape/Inactive) statuses. The independence among these categories was tested with Fisher’s exact test method.

### Regulatory regions under positive selection

The HACER database (Wang et al., 2019) was used to annotate intergenic windows in order to study potential signals of positive selection in enhancer-like regions. HACER annotates a total of 1,676,284 active enhancers (whole genome) detected by different methods (GRO-seq, PRO-seq and CAGE) in numerous cell lines and supported by different databases (VISTA, ENCODE Enhancer-like Regions, The Ensembl Regulatory Build and chromatin state segmentation by ChromHMM) which, integrated with variation data, provides a useful resource to hypothesise about the origin of non-genic signals of natural selection. In order to reduce the noise and provide a higher confidence to our intergenic signals, we have used the enhancers that at least are supported by the annotation of one database which, in the X chromosome, leave a total of 23790 active enhancers. In HACER, a given region can be annotated as an active enhancer in different cell lines, targeting the same closest gene but presenting slightly different coordinates. In order to deal with the different cell-type-specific annotations we created a “consensus” dataset of enhancers by using genomic windows. We collapsed the multiple cell-type annotations to unique enhancer coordinates when there are different overlapping enhancer regions, active in different cell lines, targeting the same gene and overlapping continuous windows. In this way we ended up with a final dataset of 1322 consensus enhancers that we used to annotate our intergenic signals. When extracting the top hits under positive selection (99.9^th^ percentile) we only took into account those enhancers that are supported by 3 or more databases in the HACER annotation, in this way we only considered high confidence enhancers that might present signals of positive selection.

### Luciferase analysis

Enhancers peaks from the top candidates were selected upon the ENCODE signals. Ancestral (A) and derived (D) haplotypes were amplified by PCR from male (*KDM6A:* NA07357 (A), NA12003 (D); *SH2D1A:* NA18501 (D)) and female (*SH2D1A:* NA18502 (A); *HUWE1*: NA18502 (A), NA18861 (D)) individuals, after checking for homozygosity, using the following primers and the KAPA high-fidelity Taq polymerase:

KDM6A (F): 5’-CATCAGAGCTCCTCTAGGCATGGGAGGGAGT-3’
KDM6A (R): 5’-TCATCTCGAGCCAGTAAGAACCTACTAGGGATCA-3’
HUWE1 (F): 5’-CATCATCTCGAGGACCAGCCACTGGGTGTAGT-3’
HUWE1 (R): 5’-TCATAAGCTTTAGGGTCCATGGTCTTCTGG-3’
SH2D1A (F): 5’-CATCATCTCGAGACAAATGTTATTGATTCCCTC-3’
SH2D1A (R): 5’-TCATAAGCTTCGACCTAAAAGAGTATA-3’

Cloning into the PGL4.10 luciferase clone was performed by using XhoI, HindIII or SacI restriction enzymes. Renilla vector was used to normalize the values as a control of transfection. Transfection into 293T cells was performed by using Lipofectamine 3000 (Thermo Fisher, L3000001), using 100 ng of luciferase and 1ng of Renilla control vector and maintained for 48 hours in OptiMEM. Cells were harvested and luciferase activity was measured using the Dual-GLO kit (Promega, E2920). Luciferase/renilla ratio calculated in 4 replicates and 2 independent experiments.

## RESULTS

We inferred recent positive selection in human X chromosomes using genomic data from 1,511 individuals of 15 populations. We conducted selection scans by applying the haplotype-based statistics iHS, iHH12 and nSL, which were designed to detect signatures of hard and soft sweeps (see Methods for details) and can be used as complementary selection tools. To assess whether a region has evolved under recent positive selection we performed coalescent simulations with *msms* (Ewing and Hermisson, 2010) to build the expected distributions under neutrality, considering human demography and the particular ascertainment bias of our data. We observed a good fit of our neutral model by comparing the observed site frequency spectrum (SFS) of the fifteen populations with their neutral simulations (Supplementary Figure 1). Small deviations in singletons are observed in some populations, but with a tight fit of alleles segregating at intermediate and high frequencies.

### Regions under putative positive selection

The per-SNP metric scores might reflect the presence of particularly homozygous regions, which could indicate the location of a selective sweep in the genome. In order to detect these signatures, the selection scores were averaged separately across sliding overlapping windows (see Methods; Supplementary Figure 2), which in most populations show distributions with a larger tail as compared with the simulations (Supplementary Figure 2A). We considered two cut-offs based on the simulated data (99^th^ and 99.9^th^) in order to extract the putatively selected windows in the empirical distributions (Supplementary Table 1).

Putative selective sweeps in regions under positive selection might present different degrees of “softness”. As noted by different authors, hard and soft sweeps are sometimes difficult to differentiate (Messer and Petrov, 2013; Schrider et al., 2015), and regions under selection might be captured by methods designed to detect both selection processes at the same time. In order to study the signature similarity in the regions under selection, we assessed the degree of overlap between the signals reported by the three metrics. Under the 99^th^ percentile in the global population, the general trend shows that iHH12 presents a similar proportion of commonly targeted regions as with iHS and nSL (~60%), while iHS targets fewer common regions as with nSL (~36%). This could be expected since iHH12 and nSL are sensitive to both hard and soft sweeps (Ferrer-Admetlla et al., 2014; Torres et al., 2018), and iHS depends on recombination rate, which might differentiate these signals from nSL signals. However, the signal overlap proves that some regions might present mixed properties of hard and soft sweeps, which could be due to the mode of selection, the degree of softness or a linked selection effect (Schrider et al., 2015).

We observed a larger proportion of signals that fall outside the simulated distribution in the African populations in the three selection tests, in comparison with non-Africans. These results are in line with previous reports which show that the number of detectable selective sweeps by haplotype-based statistics is correlated with the effective population size (Johnson and Voight, 2018; Voight et al., 2006) (Supplementary Table 1). When comparing both hard and soft selection processes we observed that soft-sweep-like signals reported by nSL and iHH12 are more abundant and widespread along the X chromosome, as was previously reported at genomic level (Messer and Petrov, 2013; Schrider et al., 2017).

The analysis reveals that high statistic values are clustered in specific spots of the X chromosome, indicating the presence of putative selective sweeps in these regions (Figure 1) (Voight et al., 2006). The distribution of signals of selective sweeps along the X chromosome is more similar between non-African than with African populations in both selection processes, indicating a common clustering of extreme signals among the different out-of-Africa populations. This was noted by Pickrell et al. (2009) and might reflect the common origin of the out-of-Africa populations and must have been acquired since leaving the African continent.

**Figure 1.**
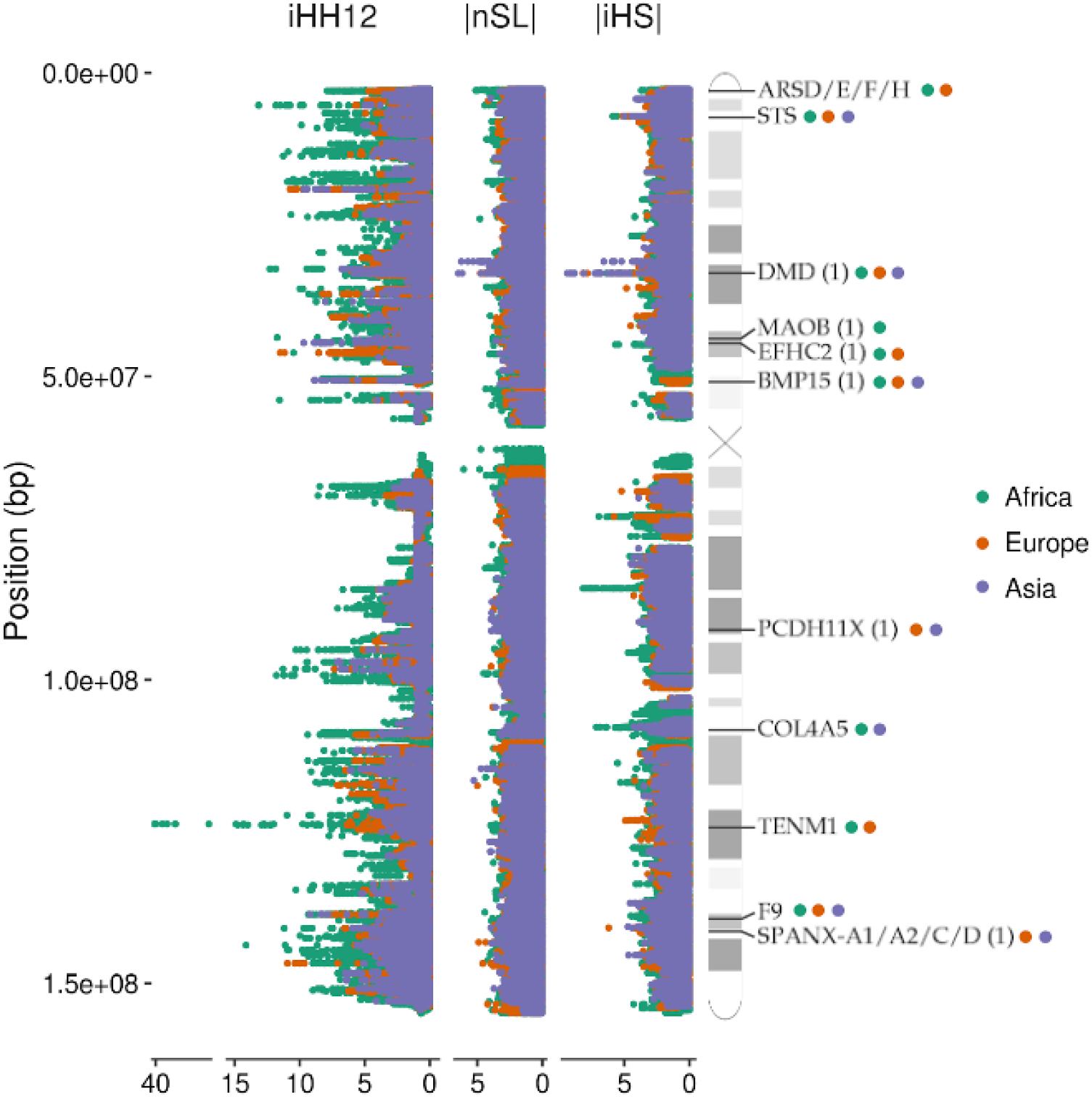
Manhattan plots of the X chromosome showing the distributions of the three selection tests used in the analysis. Some examples of genes found under selection in continental groups (99^th^; coloured circles) are shown in the ideogram. Candidates found in previous studies are indicated with (1).

### Comparison with autosomes

The unique inheritance rules of the X chromosome might generate different selection patterns as compared with the rest of the genome. In order to contrast the X chromosome signatures, we assessed selection on the autosomes of three populations of reference (Yoruba, YRI; Utah residents with northern and western European ancestry, CEU; Han Chinese, CHB) and compared the score distributions in the three haplotype-based statistics (iHS, nSL, iHH12). We see similar patterns of selective sweeps across the different populations as in the X: a higher number of outlier regions fall into the extreme tails of the autosomes in Africans (YRI) than Europeans (CEU) or Asians (CHB) (Supplementary Table 2). As seen in the X, a higher number of windows under selection are captured by the statistics nSL and iHH12 in comparison with iHS across the autosomes, probably due to the higher presence of soft-sweep-like signatures across the genome. One large difference that stands out, is that the X chromosome exhibits a consistent increase in the nSL extreme tail score distributions in non-African populations (Supplementary Figure 3). We evaluated the nSL scores in the top distribution quartile and decile, and found significant differences between the X chromosome and the pooled scores of autosomes (CEU, Kruskal Wallis: 36.04, p = 1.93e-09; CHB, Kruskal Wallis: 93.62, p = 3.81e-22). These higher selection values might be a reflection of the effect due to the haploid state in males and the smaller effective population size of the X (Veeramah et al., 2014; Johnson and Voight, 2018). However, it is difficult to associate these differences with a higher selection efficiency due to the faster-X effect, since the top 1% shows inconsistent distributions across the genome due to the presence of extreme outliers. This result might indicate that the faster-X effect is not properly captured with these selection statistics and other causes might be behind the differences seen in the distribution extreme tails.

### Gene ontology in the candidate regions

Generally, the closest gene to the estimated sweep is considered the best candidate for the target of selection. Putative selected regions were annotated as genic (protein-coding and non-protein coding; Supplementary File 1) where at least 1 bp of the window overlaps with Refseq gene coordinates. We do note that some caution is required when interpreting these results, as the strongest and widest signals are likely to span more than the target of selection.

To determine which processes are likely under selective forces, we performed a functional enrichment analysis with *Webgestalt* (Liao et al., 2019) on the top 100 putatively selected genes across all populations for the two selection regimes. There is a ubiquitous enrichment in neural-related terms in the three continental groups (Supplementary Table 3). In the two selection processes we report numerous synaptic and dendrite-related terms (e. “postsynaptic membrane” (GO:0045211), “dendrite” (GO:0030425)) with genes like *DMD, IL1RAPL1* and *GABRA3*, among others. Neuron-surface specific genes are also highly represented among the enriched terms with kinases like *CASK*, channels (*TRPC5*) and neuroligins (*NLGN4X, NLGN3*), which present their own term in numerous populations (“neurexin family protein binding” (GO:0042043)). However, for the African populations (Supplementary Table 3A) “sulfuric ester hydrolase activity” (GO:0004065) and “endoplasmic reticulum lumen” (GO:0004065) are consistently enriched non-neurological terms represented by members of the arylsulfatase family (*ARS*) and steroid sulfatase (*STS*) gene (Holmes, 2017). These genes, which are involved in hormone metabolism and are associated with X-linked diseases like chondrodysplasia punctata (Franco et al., 1995) and ichthyosis (Basler et al., 1992), present a strong signal of selection (99.9^th^) in African populations. We also observe genes consistently selected in continental groups which do not correspond with any enriched term, including reproduction-related genes, like *SPANX-A1/A2/C/D* and *SPANX-OT1* in non-African populations. These genes belong to the spermatogenesis-related gene family *SPANX-A/D*. This is a highly paralogous hominin-specific group of genes which are expressed post-meiotically in testis and some cancer types (Westbrook et al., 2006) whose members were previously reported as positively selected (Casto et al., 2010; Kouprina et al., 2004) and related to male fertility (Urizar-Arenaza et al., 2020). We observe signals of positive selection on the *BMP15* gene, related to ovarian insufficiency in women and subjected to positive selection in Hominidae clade (Ahmad et al., 2017). Other spermatogenesis-related genes (*SAGE1, SEPT6, CDK16*)and genes involved in human fertility (*ADGRG2, DIAPH2, FAM122C*) also appear in the highest scoring regions (99.9^th^) of our scans (Supplementary File 3).

### Shared sweeps in human populations

Previous reports have shown that signatures of positive selection are often shared between different human populations (Johnson and Voight, 2018). Common evolutionary trajectories might generate similar selective pressures which leave shared signatures of positive selection. These common patterns might reveal important traits that were crucial in the adaptation of ancestral populations. To that end, we assessed the degree of sharing of the candidate regions under putative positive selection. We considered the 99^th^ percentile candidates in the three selection tests and identified those regions whose genomic coordinates overlap across multiple populations. We found that 41% of the selective sweeps are unique to a specific population, 38% are shared between populations of the same continental group and 20% are shared across different continents. These results are in line with previously reported selection patterns (Johnson and Voight, 2018): common sweep events are more frequent between closely related populations, and cross-continental sweeps are rarer and more likely to result from common selective pressures and older processes of positive selection.

Among the cross-continental selected regions we found that one of the most commonly shared falls within the *DMD* (dystrophin) gene. This is the largest gene in the human genome and is involved in the stabilization of the sarcolemma and synaptic transmission. We found multiple signatures of hard and soft sweeps across the 15 included populations, which together span a region which reaches up to ~2Mb (Supplementary Figure 4A). The variable length of this sweeping region might indicate that multiple selection events took place in the three continental groups, which generated different patterns that suit the two selection processes. Positive selection signals were previously reported in several components of the dystrophin protein complex (*DPC*) (Williamson et al., 2007) in non-African populations and in *DMD* in Africans (Casto et al., 2010). Our *DMD* results are complementary to these previous studies and validate evidence for adaptations in neurological and muscle-related phenotypes in other populations.

Another globally shared region overlaps the *F9* gene, which encodes the coagulation factor protein FIX and is involved in Hemophilia B. In this case, the *F9* region harbours windows under positive selection in the 99.9^th^ percentile reported by iHH12, which reflects a sweeping region that spans up to ~50kb (Supplementary Figure 4B). A previous study reported coagulation factors underwent positive selection in different clades (Rallapalli et al., 2014), which might be a consequence of selective pressures due to the direct relationship with the immune system and host-pathogen interactions. Although the FIX factor has not been identified as related to any selective pressure to date, it might be under recent positive selection in human populations due to its role in the coagulation system as the first line of defence against pathogens.

### TENM1 gene

The most extreme signals in the analysis are reported by iHH12, reaching in some cases values between 10 and 15 in African populations (>99.97%). Patterns of soft and incomplete hard sweeps might be a side effect of linked regions targeted by complete hard sweeps, referred to as the “soft sweep shoulder” (Schrider et al., 2015). A possible example of this is seen in the *TENM1* gene, which is the highest scoring region in the chromosome with an iHH12 signal composed of two high peaks (Figures. 2A,B). This gene is involved in neural development and is specifically determinant for the synapse organization of the olfactory system. In African populations this region exhibits a peak value of iHH12 > 40, while in non-African populations is hardly captured by iHH12 due to an excess of low minor allele frequency variants (MAF < 0.05), which are filtered out by *selscan*. iHS and nSL outlier windows are also found within this region, suggesting the presence of haplotype patterns which fit with both soft and hard sweep signatures. In order to elucidate the haplotype structure of this region, we inferred clusters of similar haplotypes with *fastPHASE* (Scheet and Stephens, 2006) on representative populations of the three continental groups (CEU, CHB and YRI). Figure 2C shows different long haplotypes at high frequency with the main presence of two highly homozygous clusters overlapping the iHH12 peaks, either in African or non-African populations. This pattern is expected in regions that underwent selection processes and left long, unbroken haplotypes where no recombination events occurred. The two main clusters span ~300kb of the *TENM1* gene and their location suggests that an ancient strong selection event took place in this region before the population split in the out-of-Africa event. For confirmation, we calculated the Tajima’s D statistic, which was designed to detect ancient complete sweeps (Pybus et al., 2015), in all the populations. Figure 2B depicts the spanning region which presents an ancient complete hard sweep with windows that reach a Tajima’s D ≤ −2 (1% extreme). This suggests that, despite not observing iHH12 signals in non-African populations, the underlying haplotype pattern reflects a signature of positive selection that includes the global population. No clear phenotype could be associated with this signal, however recent evidence indicates mutations in *TENM1* are linked with congenital general anosmia (Alkelai et al., 2016), suggesting the potential for olfactory adaptations. Previous studies have shown the importance of the olfactory system in the evolution of *Homo sapiens* (Hoover, 2010), olfactory receptors were subjected to non-neutral selection (Hoover, 2015) which accounts for population-specific phenotypic variability (Trimmer et al., 2019). This evidence suggests that olfactory receptors, and the associated neural system, might be subjected to important adaptive processes in human evolutionary history.

**Figure 2.**
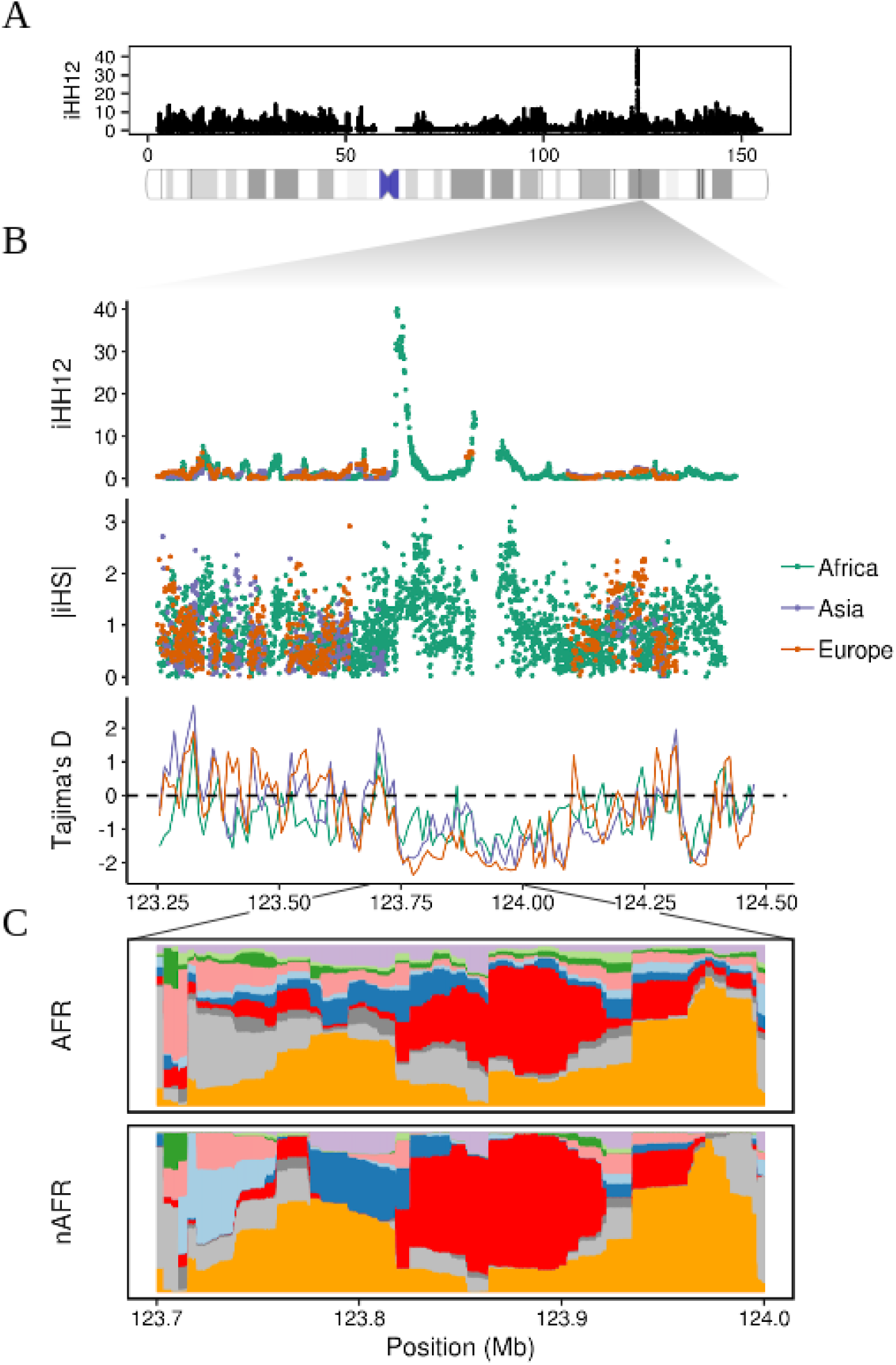
Putative positive selection signal on the *TENM1* gene. **(A)** Whole chromosome iHH12 scores in the global sample. **(B)** Manhattan plot showing the iHH12, iHS and Tajima’s D scores on the *TENM1* gene region. **(C)** Clusters of highly similar haplotypes (in red and orange) estimated by *fastPHASE* were found in African (AFR) and non-African populations (nAFR). The different colouring represents changes in the haplotypic composition through the region, where each row represents a haplotype and each column a SNP.

### Selection of X-inactivation escape genes

The incomplete inactivation of some genes, during the process of gene dosage compensation in females, might expose these escapees to sex-especific adaptive processes due its biased expression. We wanted to investigate whether patterns of positive selection could be detected amongst the genes that escape from the X chromosome inactivation. We obtained the X chromosome inactivation (XCI) status in the combined set of populations from Tukiainen et al. (2017). We considered 59 genes as “escape” and 381 genes as “inactive”, keeping only genes with the strongest support. We constructed contingency tables based on these categories performing Fisher’s exact test of independence between selection and XCI status for different extreme tail thresholds of the selection tests. We found that genes that escape from the X-inactivation had a higher probability of being targeted by positive selection according to two of the tests. This trend is significant for iHS, is marginally significant for iHH12 and does not reach significance for nSL (Supplementary Tables 4A,B). Notably, escape genes under positive selection had similar proportions from iHS and iHH12 (19% and 20%, respectively; Supplementary Table 4A), however only reached 11% for nSL. This may suggest that escape genes are more likely to be targeted by selection processes that leave signatures with a degree of “softness” closer to hard sweeps rather than soft sweeps.

Supplementary Table 4C lists the genes under selection that escape inactivation. On this list, we found enrichment in sulfuric ester hydrolase activity (GO:0008484), due to the sulfatase group of genes. Among these top candidates, we found four members of the *ARS* family. Three of these members participate in bone and cartilage matrix composition during development (*ARSE*, *ARSD*, *ARSF*). These genes are associated with the X-linked Chondrodysplasia Punctata, a syndrome that affects almost exclusively females, and is characterized by abnormal embryo development, including skeletal malformations, skin abnormalities and cataracts (Franco et al., 1995).

The *STS* gene, also escaping inactivation, presents another highly shared sweeping region among populations (iHS 99.9^th^ percentile in African populations and 99^th^ percentile in Europeans and Asians). It is associated with the X-linked Ichthyosis, a syndrome caused by a placental steroid hormone deficiency and is characterized by skin and eye abnormalities (Basler et al., 1992). This gene was reported to be one of the top female-biased genes differentially escaping inactivation in Yoruba (YRI) (Johnston et al., 2008). As hypothesized by Tukiainen et al. (2017) most of the escape genes reported as under selection show female-biased expression, suggesting these genes might be involved in some adaptive trait in females.

### Functional non-coding regions under positive selection

Previous studies have reported numerous signatures of positive selection with an unknown coding genic cause. This might be accounted by a high false positive rate in genomic scans but also by the presence of signatures in non-genic regions, suggesting that many true signals are located in non-coding, potentially regulatory elements (Fraser, 2013; Enard et al., 2014).

In order to identify the strongest and most interesting candidates of positive selection on the X chromosome, we evaluated the signals in the 99.9^th^ percentile and attempted to pinpoint the target of selection within each signal by annotating SNPs with *ANNOVAR* (Wang et al., 2010). A large portion of single nucleotide polymorphisms (SNPs) over the 1% per-SNP score extreme tail are intergenic, in addition, a large fraction fall within intronic regions for all statistics (iHS: 0.29, iHH12: 0.32, nSL: 0.2), with few in exons or untranslated regions. Combined Annotation Dependent Depletion (CADD) scores (Kircher et al., 2014) were used to identify functional variants according to their deleteriousness (see Methods). After filtering by functionality (CADD ≥ 10), the majority of the variants were excluded, however, the SNP composition remained higher in intergenic regions (Supplementary Table 5), with an average prevalence in signals reported by iHH12 and nSL in non-African populations (Africa: ~0.62, Europe: ~0.72, Asia: ~0.9). These results suggest that there is an excess in signals driven by intergenic SNPs that fall in non-annotated and potentially regulatory regions.

Several intergenic regions are under positive selection in the different continental groups (Supplementary Table 1). In order to assess the functional impact of these signals, we explored the overlap of the extreme SNPs within the 99.9^th^ percentile windows with RegulomeDB (Boyle et al., 2012) annotated elements. The combined signals across all populations had higher proportions of SNPs within an ENCODE element (iHS: 19.1%, iHH12: 26.3%, nSL: 13%) compared to the whole chromosome (5.5%). This enrichment is more prevalent for iHH12 signals, which may be due to its power to detect both hard and soft sweep signatures. This finding shows, as expected, intergenic regions under putative positive selection are enriched in functional elements and likely points to selection of regulatory processes.

Intergenic signals cluster around genic regions, suggesting a regulatory function influencing surrounding genes. Under the 99^th^ percentile, we found instances of genic windows that overlap genic and intergenic SNPs, this is more prevalent in iHH12 and nSL statistics (iHS: 2%, iHH12: 5.7%, nSL: 4.4%) across all populations. Since regulatory elements are expected to be found in the extremes and within coding regions, we used the RegulomeDB annotation to associate the signal of putative selection with any potential regulatory function. In these overlapping regions we found that the odds of intergenic SNPs overlapping a functional element is higher than genic SNPs (Supplementary Table 6A) according to iHH12 and nSL, moreover when considering extreme SNPs (99.9^th^) these values reflected a much higher dominance of functional intergenic SNPs in these tests (Supplementary Table 6B). These findings indicate that the overlapping genic windows under selection are more enriched in regulatory elements in their intergenic portion, something that points to the presence of sweeps in regulatory elements.

This evidence suggests, as previously noted, amino acid changes may play a less important role in recent adaptation and that regulatory changes may drive a more important part of adaptation events in recent human evolution (Fraser, 2013; Enard et al., 2014; Grossman et al., 2013).

### Enhancer-like signatures under positive selection

In order to analyse in more detail the regulatory roles of the regions under putative positive selection, we intersected the intergenic windows in the extreme tails with the enhancer coordinates described in the Human Active Enhancer to interpret Regulatory variants (HACER) database (Wang et al., 2019). Supplementary Table 7 shows the overlapping/non-overlapping windows with enhancer regions (in any cell line) in the 99^th^ percentile extreme tail. As the table shows, the intergenic regions under positive selection are more probable (odds ratio values) to present overlapping enhancers in the case of iHH12.

In several cases these enhancers were located close to genes also reported as positively selected in the analysis. We wanted to determine if this pattern is a by-product of the selection in adjacent regions by genetic linkage (hitchhiking effect), or due to independent selection processes on both elements, the enhancers and their target genes. In order to deal with the different cell-type-specific enhancers described in HACER we created a consensus enhancer dataset (see Methods) with unique coordinates. We pooled all the populations and selection tests in order to maximize the statistical power of our analysis. A Chi-squared test shows the dependency between the selection of the enhancers and their target genes (p-value = 0.0021). However, despite the dependency between these two variables we observe a higher probability of both elements, the enhancer and its closest gene, as being under positive selection (YY category) and not being under positive selection (NN category) than expected by chance (Supplementary Tables 8A,B). We compared the mean distances between the selected/non-selected enhancers and their selected/non-selected closest genes. These distances do not seem to support the physical genetic linkage as a possible explanation of this association. It must be taken into account that the reported distances are sometimes too large (~2.5Mb) to be the reason for selection by hitchhiking of both elements. Therefore, the YY set of enhancers and target genes must be regions that are jointly swept by hitchhiking (most of them) combined with few regions that are selected by independent processes. This suggests that selective pressures might affect some genes and their regulatory elements in a coordinated way, modifying not only their coding sequence but also their expression level.

Next, we wanted to study the potential origin of some of the most extreme intergenic signals and the regulatory effect of the sweeping haplotypes in the different populations. We focused on the highest scoring candidate enhancers (99.9^th^) and their closest genes (Supplementary Table 9). Among these candidates, we found at X:73,135,561-73,145,161 an iHS African-shared extreme signal that overlaps an enhancer (Supplementary Figure 6) located in the XIC region (X-inactivation center) and whose closest gene is *JPX*. This region is active in five different cell lines according to HACER (H1, HUVEC, HCT116, AC16, REH) and is supported by three databases (Ensembl Regulatory Build, ENCODE Enhancer-like Regions and ChromHMM). The gene *JPX* (~23kb away) is an activator of the lncRNA *XIST*, which is involved in the X chromosome inactivation. Among the potential causal variants of this signal, the SNP rs112977454 reported as expression quantitative trait loci (eQTL) by the Genotype-Tissue Expression (GTEx) project, is the most likely candidate. In addition, this eQTL has a CADD score of 9.018, close to the 1% pathogenicity threshold (CADD = 10) used by Kircher et al. (2014), and an average derived allele frequency (DAF) of 17% in African populations, while is absent from the rest of populations. This eQTL is also found overlapping a transcription factor binding site (TFBS) in the HUVEC cell line, which targets the *JPX* gene through the transcription factors *FOS, GATA2, JUN* and *POLR2A*. No specific phenotype is associated with this variant; however, these results suggest that its segregation in African populations might influence the transcription factor binding and affect the regulation of the *JPX* gene.

### Functional analysis of enhancers under positive selection

In order to explore the potential regulatory effect behind the selection processes in the candidate enhancers (Supplementary Table 9), we compared the regulatory activity of the putative haplotype under selection with that of its ancestral sequence. To perform this task, we analyzed the changes in the expression of the reporter gene luciferase under regulation of the two ancestral and derived haplotypes in some of these enhancers. This method allows us to test all the potential causal variants independently on the possibility of testing a passenger variant (not causal) of the sweep. We tested the enhancer regions targeting the genes *HUWE1, KDM6A* and *SH2D1A* (Figures 3A,B,C), which also harbor signals of positive selection in their sequences. These genes are implicated in intellectual dissability (*HUWE1*) (Giles and Grill, 2020) and the Duncan disease (*SH2D1A*) (Sumegi et al., 2002), and, in the case of *KDM6A*, this gene is reported as X-inactivation escapee by Tukiainen et al. (2017), which makes it susceptible to participate in sex-specific processes (Dunford et al., 2017, Itoh et al., 2019). In all these cases, the enhancer region overlaps with more than one potential causal SNPs, located almost all of them in the 99^th^ percentile of the selected populations. Ancestral and derived haplotypes of the candidate enhancers were obtained from males of the relevant population under selection and subsequently cloned in a luciferase-reporter vector. Upon transfection in 293T cells, significantly differential luciferase activity amongst the ancestral and derived haplotypes for *HUWE1* and *KDM6A* enhancers was observed, showing a clear distinction of the regulatory activity between these two haplotypes (Figure 3D). Yet this analysis did not show differential activity between the ancestral and derived form of the *SH2D1A* enhancer. Although no specific phenotypes were able to be assigned to the selection of these regions, our data suggest that positive selection has contributed to the adaptation of different human populations by differentially regulating the expression of certain genes. Further studies will be needed to understand the phenotypic consequences of such adaptations.

**Figure 3.**
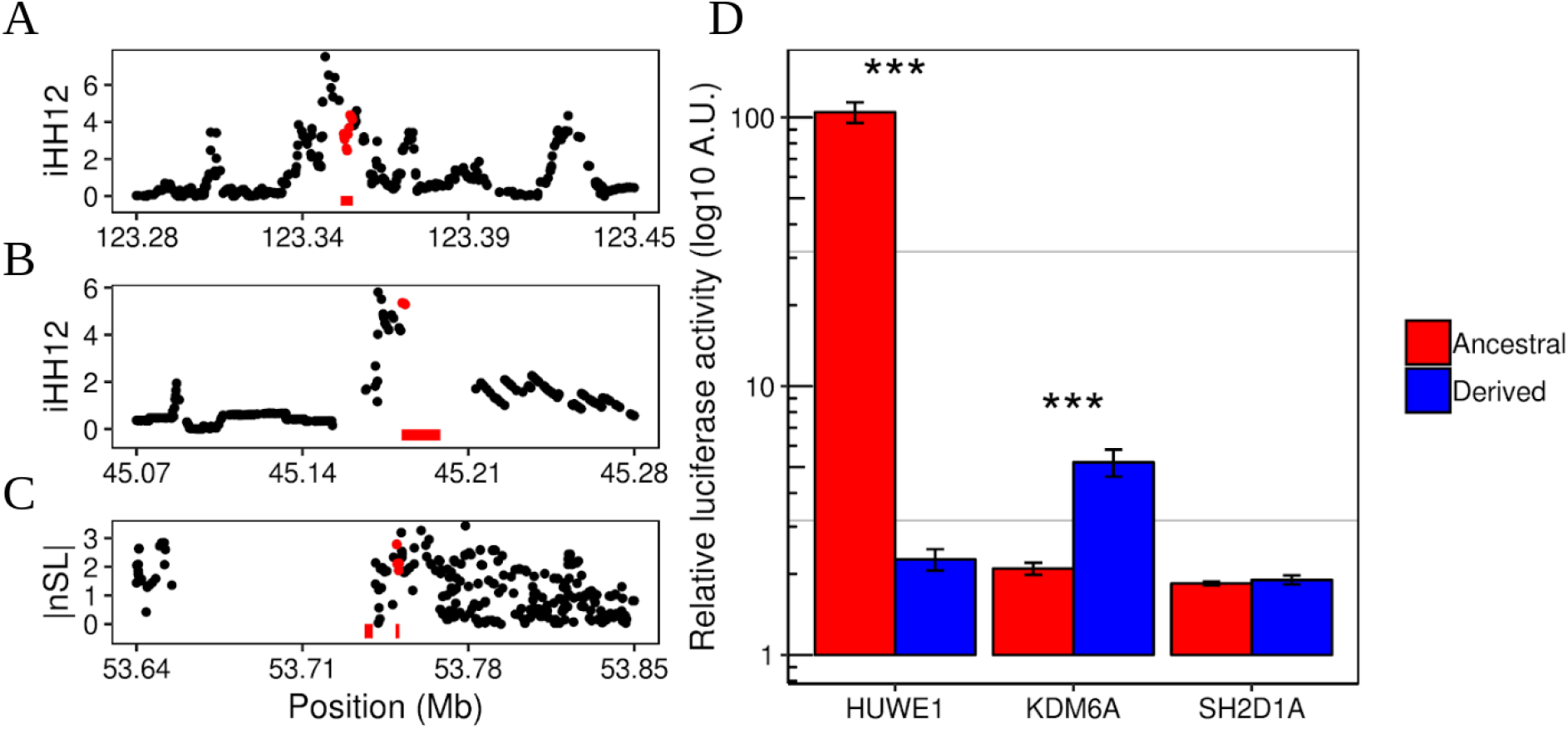
Candidate enhancers under putative positive selection. Manhattan plots show the selection scores overlapping the enhancer coordinates (bottom red bars) targeting *SH2D1A* **(A)**, *KDM6A* **(B)** and *HUWE1* **(C)** genes in YRI, CEU and YRI populations, respectively. Although *HUWE1* appears under positive selection in Gambians (GWD) (Table S9) YRI individuals were used in the luciferase assay instead, since the signal is also present in this population at 99^th^ percentile. Red dots correspond to enhancer overlapping SNPs. **(D)** Relative luciferase activity comparisons between the ancestral and derived haplotypes in each of the candidate enhancers. Significant differential activities are seen in *HUWE1* (p-value = 5.75e10^-8^) and *KDM6A* (p-value = 0.004) enhancers.

## DISCUSSION

In this analysis, we report a comprehensive analysis of recent positive selection in the X chromosome of 15 non-admixed sub-Saharan African, European and East Asian populations. We have focused on the spectrum of signatures captured by the selection statistics iHS, iHH12 and nSL, which are based on the detection of extended long haplotypes at moderately high and intermediate frequencies (hard and soft sweeps). These three statistics present different approaches and statistical power to detect the different modes of selective sweeps. However, in some cases, the similarity between the haplotypic patterns behind hard and soft sweep signals might lead to the simultaneous detection of the same selected region by these three methods. Results indicate Sub-Saharan African populations have a higher proportion of windows that fall outside the extreme simulation thresholds in comparison with Europeans and Asians. This is directly related to the effect of haplotype-based statistics, in which the number of detectable windows under selection is correlated with the effective population size (Johnson and Voight, 2018; Voight et al., 2006). In contrast with iHS, a higher amount of soft sweep-like signatures is presumably captured by nSL and iHH12 statistics. This was previously noted by authors who claimed that regions targeted by hard sweeps are much less common than soft sweeps (Messer and Petrov, 2013; Schrider and Kern, 2017). Subtle changes of frequency in multiple loci might be also behind numerous quantitative adaptations that would require a more profound and comprehensive analysis than the one conferred by the “sweep” vision (Höllinger et al., 2019). Therefore, it is more likely that genomes, and the human X chromosome in this case, are populated by a greater number of signatures with different degrees of “softness” that are misclassified or overlooked by most selection statistics.

The faster-X effect is believed to act on the X chromosome when the hemizygous state leads to a complete penetrance of mutations, allowing for a quicker and stronger adaptive process. Differences between autosomes and the X chromosome are seen for the nSL statistic in non-African populations, which might suggest some kind of effect that generates the skewed distributions. However, these differences could not easily be associated with the faster-X effect, due to the inconsistencies found in the top 1%. However, as previously noted by Arbiza et al. (2014), natural selection seems to be a more powerful force in the sexual chromosome than in autosomes, which might explain differences in X/Autosome diversity in human populations. Particular selection events and sex-biased processes might leave specific pronounced signatures in the X chromosome, like we report in this paper. Nevertheless, despite accounting for demography and different mutation rates in our simulations, selection is not the only factor that could be invoked to explain the differences in haplotype diversity.

We report signals of recent positive selection in particular regions of the X chromosome. The difficulty of identifying clear signals from particular selection processes relies on the mixed properties of most signatures. In our scan most of the observed signals are captured by more than one statistic. One of the most remarkable cases of selection in our analysis is the *TENM1* gene. This gene harbours a region of ~300kb (Figure 2) with selection signals that indicate the presence of a haplotype pattern indicating an old and strong event of positive selection before the human populations split. Moreover, the haplotype clusters inferred by *fastPHASE* show a clear predominance of two types of sequences that might derive from a whole unique sweeping haplotype that could be broken by recombination in this hotspot region. Although the role of *TENM1* selection might be linked to recent changes of the olfactory system, the origin of the haplotype patterns seen in our analysis could have more general implications for neural development. Genic regions under putative positive selection seem to be dominated by genes involved in neural development enriched processes. This is widely reported by the three tests used and appear globally distributed in the three continental groups. These findings fit the general picture of previous evolutionary studies which describe the role of neural genes in human recent history (Wei et al., 2019).

Commonalities with previous studies reinforce evidence of X-linked selection in human populations. Despite differences in the approach, we found complementary results. Indeed, the great diversity between populations in our study has confirmed previously described signals, like selection in *DMD* or reproduction-related genes like the *SPANX* family, and expanded the findings in new populations and genes. It is of interest to remark on the case of the *SPANX* members and other reproduction-related genes reported above. It was previously mentioned the potential importance of fertility-related genes in recent human history (Ramm et al., 2014; Hart et al., 2018). The *SPANX* members, like other cancer-testis (CT) genes (*MAGE* family in the 99^th^ percentile), are known to be under rapid evolution and appear to be subjected to positive selection affecting their coding sequences (Kouprina et al., 2004). Previous reports found members of the spermatogenesis-related family *SPATA* to be under recent positive selection and suggest that testis-enriched genes are the target of population-specific selection (Schrider and Kern, 2017; Schaschl et al., 2020). Other studies report specific ampliconic gene-enriched regions in humans and other primates targeted by strong selective sweeps, where meiotic drive and sperm competition seem to be a potential explanation (Dutheil et al., 2015; Nam et al., 2015). Although an important number of previously reported genes under selection have been captured in our scan, it is important to note that a high false discovery rate is expected from this “hypothesis free” approach. Nonetheless, despite the likely presence of false positives, our findings are in line with previous evidence and supports the importance of reproduction and male fertility in recent human evolutionary history.

The gene dosage compensation of the X chromosome occurs in females by the random inactivation of one of the copies during the early stages of embryogenesis. However, this process of transcriptional silencing is not complete for all the genes. Evidence suggests that around 23% of the X-linked genes “escape” inactivation and express both chromosomal copies. Most of these genes are located in the pseudoautosomal region 1 (PAR1) and only a small fraction is distributed along the non-pseudoautosomal region (nPAR) (Balaton et al., 2015; Tukiainen et al., 2017) analysed in this study. Overall, our analysis shows an enrichment of genes under selection which escape X-inactivation mainly driven by hard-sweep-like signatures. These genes were previously described as being under purifying selection (Park et al., 2010), however, no evidence for positive selection has been reported until now. Although one could argue that background selection might be behind such a pattern, a recent study has shown that this kind of selection is not expected to mimic the signatures left by selective sweeps (Schrider, 2020). Therefore, these X-linked escape genes are expression-biased between sexes and might be responsible for sexual dimorphic traits, likely producing phenotypic diversity which has been adaptive in females during human evolution. However, more specific analyses on escape genes are needed in order to establish a phenotypic cause for such potential adaptation.

A large fraction of regions under selection have no annotations. We report significant evidence of intergenic regions with high selection scores in the three selection tests, reflecting the presence of signatures that fit the two selection processes we consider in this analysis. Enrichment in the regulatory elements annotated by RegulomeDB is seen globally in the two selection processes, with a higher prevalence in regions exhibiting soft sweep-like signatures (iHH12 and nSL signals). Sometimes genic regions might be affected by the selection of the surrounding intergenic regions that harbour regulatory elements. In our analysis a fraction of selected windows classified as genic have intergenic portions that exhibit a dominance of highly scored SNPs that overlap a functional non-genic element reported by RegulomeDB.

A recent analysis of selection in enhancers revealed that approximately 5.90% of the enhancers studied in different tissues present signatures compatible with recent positive selection events (Moon et al., 2019). Other cases of selection in enhancers have shown how a SNP subjected to positive selection is able to modify the regulatory activity of the region in a population specific manner (Nakayama et al., 2017). Having this in mind we used the HACER database to study in more detail the potential role of selection in active human enhancers. We show several cases of reported enhancers under selection whose closest gene (also considered target gene) is under putative positive selection in our analysis. This result might reflect a linkage effect between these two elements; however, we suggest that in some cases this is an indication of concurrent selection of the gene and the regulatory region. We report specific cases of putative positive selection signals in enhancers that might drive population-specific regulatory changes. African populations had a highly scoring hard sweep-like signature in an enhancer located in the XIC region. Among the top SNPs we find rs112977454 (99.96^th^ percentile) as an eQTL highly segregated in Africans which might affect the binding of transcription factors that regulate the expression of the lncRNA *JPX*. This gene is a key participant in the X chromosome inactivation as it promotes the expression of *XIST* (Tian et al., 2010), which finally silences the transcription by coating the chromosome into the Barr body. This is an interesting candidate since it might affect the expression patterns of genes that escape from the X-inactivation and thus play a role in the potential adaptations of dimorphic traits hypothesized before.

In order to reveal the potential regulatory effect of our enhancers under selection, we performed luciferase-based assays on three of our top candidates that met the requirements to be cloned. *HUWE1* and *KDM6A* enhancers exhibit a significant difference in the luciferase activity between the two most differentiated haplotypes. This effect clearly suggests a differential regulation of these genes which might fit with the idea of population-specific selection processes. The case of *KDM6A* is rather remarkable since it has been associated with female-specific traits where its ability to escape from the X-inactivation plays a significant role. The biallelic expression of this gene seems to confer a protective effect in females in a wide range of cancer types, where males are more exposed due to their hemizygous state (Dunford et al., 2017). The same overexpression of *KDM6A* appears to be involved with sex differences in autoimmune disease susceptibility, contributing to a higher incidence of multiple sclerosis in females (Itoh et al., 2019). Although we were not able to make a direct association between our selection signals and these phenotypes, the evident effect of selection in these enhancers and the potential role of adaptations in escape genes suggest that selection might be behind secondary processes that affect women and men in different ways. As for other genomic scans, the power to detect regions under positive selection in our analysis might leave behind more complete patterns that explain in a more comprehensive way the potential adaptations presented here. This, together with the inherent difficulty of identifying the precise target of natural selection, make this type of analysis a challenging aspect in the study of evolution.

## List of abbreviations

CADD: Combined Annotation Dependent Depletion
DAF: Derived Allele Frequency
EHH: Extended Haplotype Homozygosity
eQTL: Expression Quantitative Trait Loci
GO: Gene Ontology
HACER: Human Active Enhancer to interpret Regulatory variants database
HMM: Hidden Markov Model
iHH12: Integrated Haplotype Homozygosity pooled test
iHS: Integrated Haplotype Score test
Kb: Kilobases
LD: Linkage disequilibrium
MAF: Minor Allele Frequency
Mb: Megabases
nPAR: Non-pseudoautosomal region
nSL: Number of segregating sites by length test
OEA: Overrepresentation Enrichment Analysis
PAR: Pseudoautosomal region
SFS: Site Frequency Spectrum
SNP: Single Nucleotide Polymorphism
TFBS: Transcription Factor Binding Site
XCI: X Chromosome Inactivation
XIC: X-inactivation center

## Authors’ contributions

JB, HL conceived the study. PV-M, SA and HL analysed and interpreted the data. PV-M wrote the manuscript. JN, HL, SA, JB revised the manuscript. All authors approved the final manuscript.

## Acknowledgments

The authors would like to thank the contribution of Andrea Martí Sarrias in the performance of the luciferase assays of enhancers under positive selection.

## SUPPLEMENTARY MATERIAL

*See supplementary figures and tables in separated files. Titles and captions are provided below*.

### Supplementary Figures (.pdf)

**Supplementary Figure 1.** Comparison of site frequency spectrums between empirical and simulated data across all populations. Fixed sites have been pruned.

**Supplementary Figure 2.** A) iHS, iHH12 and nSL distributions (dashed lines as simulated scores) and B) QQ plots of the window-based score distributions in the three geographical groups (Sub-Saharan Africa, Europe, Asia). The QQ plots indicate an overall agreement between the observed and simulated scores. An enrichment of high values in some groups (iHH12 ~ 30) are due to the presence of extreme outliers in the empirical distribution (≥99%), this can be seen in the density plots by the long tail towards positive values.

**Supplementary Figure 3.** Comparison between nSL extreme tail distributions of autosomes and X chromosome.

**Supplementary Figure 4.** Sweeping regions under putative positive selection in the three continental groups in the dystrophin gene (*DMD;* A) and coagulation factor 9 (*F9;* B). The reported sweeps are a result of merging the overlapping windows under positive selection in the 99^th^ in all the statistics used (iHS, iHH12 and nSL).

**Supplementary Figure 5.** Manhattan plot showing the putative positive selection signal in African populations reported by iHS in the enhancer located at ~23kb from the *JPX* gene. Marked in red is the SNP rs112977454 reported as eQTL by GTEx. Colour bars at the bottom represent the active enhancers in five different cell lines. Legend shows the window-based score where the SNPs belong to.

### Supplementary Tables (.pdf)

**Supplementary Table 1.** Windows under putative positive selection in the extreme simulated 99^th^ and 99.9^th^ percentiles across the 15 populations under study and the three selection statistics accounting for hard and soft sweeps (iHS, iHH12 and nSL).

**Supplementary Table 2.** Windows under putative positive selection in the extreme simulated 99^th^ percentiles in the human autosomes of the three populations of reference (YRI, CEU and CHB) across the three selection statistics accounting for hard and soft sweeps (iHS, iHH12 and nSL).

**Supplementary Table 3A.** Significant GO terms of the top 100 genes across all the Sub-saharan African populations in the three selection tests used in the analysis. We consider FDR < 0.05 as significant. In the table, we present the population ID, the tests where the term is reported as significant, the GO term ID, the term description and the corrected FDR value.

**Supplementary Table 3B.** Significant GO terms of the top 100 genes across all the European populations in the three selection tests used in the analysis. We consider FDR < 0.05 as significant. In the table, we present the population ID, the tests where the term is reported as significant, the GO term ID, the term description and the corrected FDR value.

**Supplementary Table 3C.** Significant GO terms of the top 100 genes across all the Asian populations in the three selection tests used in the analysis. We consider FDR < 0.05 as significant. In the table, we present the population ID, the tests where the term is reported as significant, the GO term ID, the term description and the corrected FDR value.

**Supplementary Table 4A.** Contingency tables of escape genes under selection reported by the three selection statistics across three extreme percentiles (95^th^, 99^th^ and 99.9^th^). Two categories were used: escape/inactive and selected/non-selected.

**Supplementary Table 4B.** Fisher’s tests applied to the contingency tables. iHS reports significant p-values across the three extreme percentiles with increasing odds ratios (OR). iHH12 and nSL do not show significant enrichment in escape genes, however the odds are in line with those in iHS in five out of the six comparisons, suggesting the presence of selection but with lack of significance probably due to a sample effect.

**Supplementary Table 4C.** Escape genes reported by iHS as being under positive selection in each continental group across the extreme percentiles.

**Supplementary Table 5.** SNPs with a selection score within the 1% extreme and with CADD score ? 10 in the 99.9^th^ percentile across all populations (Intergenic (Int), Intronic (I), Exonic (E), Downstream (D)

**Supplementary Table 6A.** RegulomeDB annotation of the 99^th^ percentile genic windows with intergenic overlap and the odds ratio (OR) between genic and intergenic SNPs within functional elements. All populations are considered.

**Supplementary Table 6B.** RegulomeDB annotation of the 99^th^ percentile genic windows with intergenic overlap when considering extreme scoring SNPs (per-SNP 1% extreme tail). iHH12 and nSL show a significant OR increment in comparison with iHS

**Supplementary Table 7.** Overlapping and non-overlapping intergenic windows under putative positive selection on enhancer regions reported by HACER in any cell line (see Methods) across the three continental groups. Odds ratio (OR) of intergenic and overlapping windows shows a significant increment mainly in iHH12 across all populations.

**Supplementary Table 8.** Contingency tables of both observed and expected pairs of enhancer/target-gene in the following categories: Selected enhancer and selected gene (YY), Selected enhancer and non-selected gene (YN), Non-selected enhancer and selected gene (NY), Non-selected enhancer and non-selected gene (NN). A Chi square test is applied to study the dependency of both variables (Chi sqr value = 9.44; p-value = 0.0021).

**Supplementary Table 9.** Top enhancer regions under putative positive selection (99.9^th^ percentile). The genes marked as “**” are found under selection in sequence in the 99.9^th^ percentile and in the same continental group, the genes marked as “*” are found under selection as well but in a different continental group.

**Supplementary File 1 (.xlsx) : Supplementary gene tables.** List of genes found under putative positive selection in all populations in 99th and 99.9th percentiles.

