## Supplementary Figures for "Chromosome X-wide analysis of positive selection in human populations: from common and private signals to selection impact on inactivated genes and enhancers-like signatures"

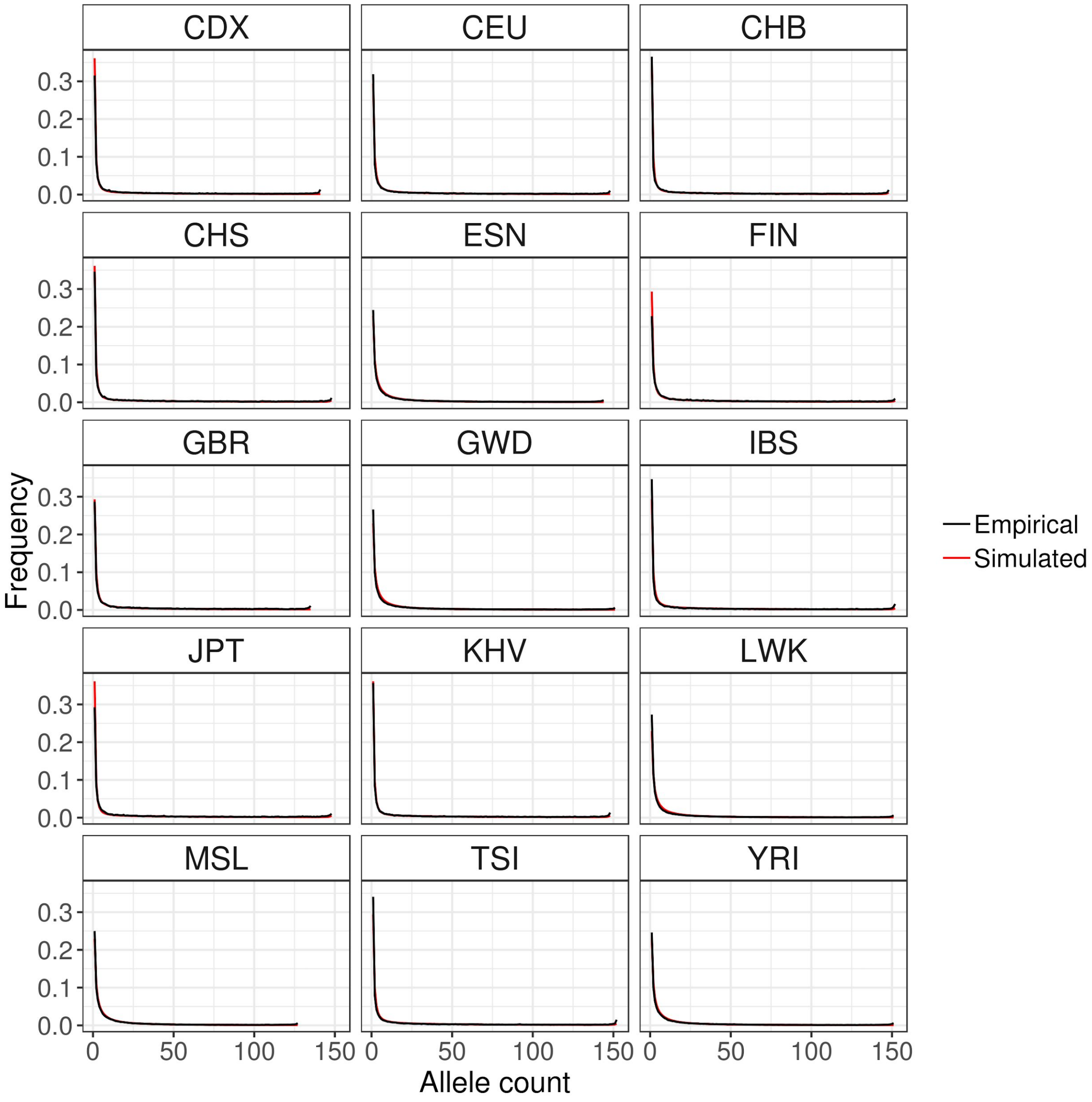


**Supplementary Figure 1.** Comparison of site frequency spectrums between empirical and simulated data across all populations. Fixed sites have been pruned.

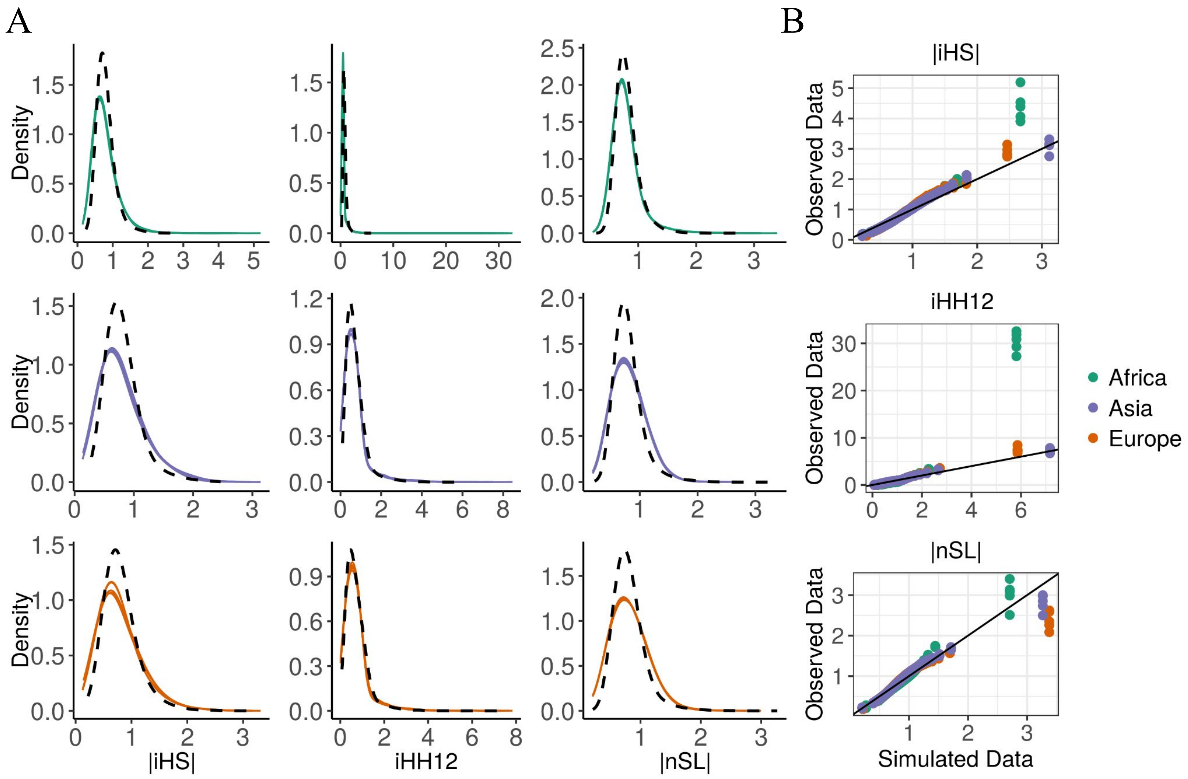


**Supplementary Figure 2.** A) iHS, iHH12 and nSL distributions (dashed lines as simulated scores) and B) QQ plots of the window-based score distributions in the three geographical groups (Sub-Saharan Africa, Europe, Asia). The QQ plots indicate an overall agreement between the observed and simulated scores. An enrichment of high values in some groups (iHH12 ~ 30) are due to the presence of extreme outliers in the empirical distribution (≥99%), this can be seen in the density plots by the long tail towards positive values.

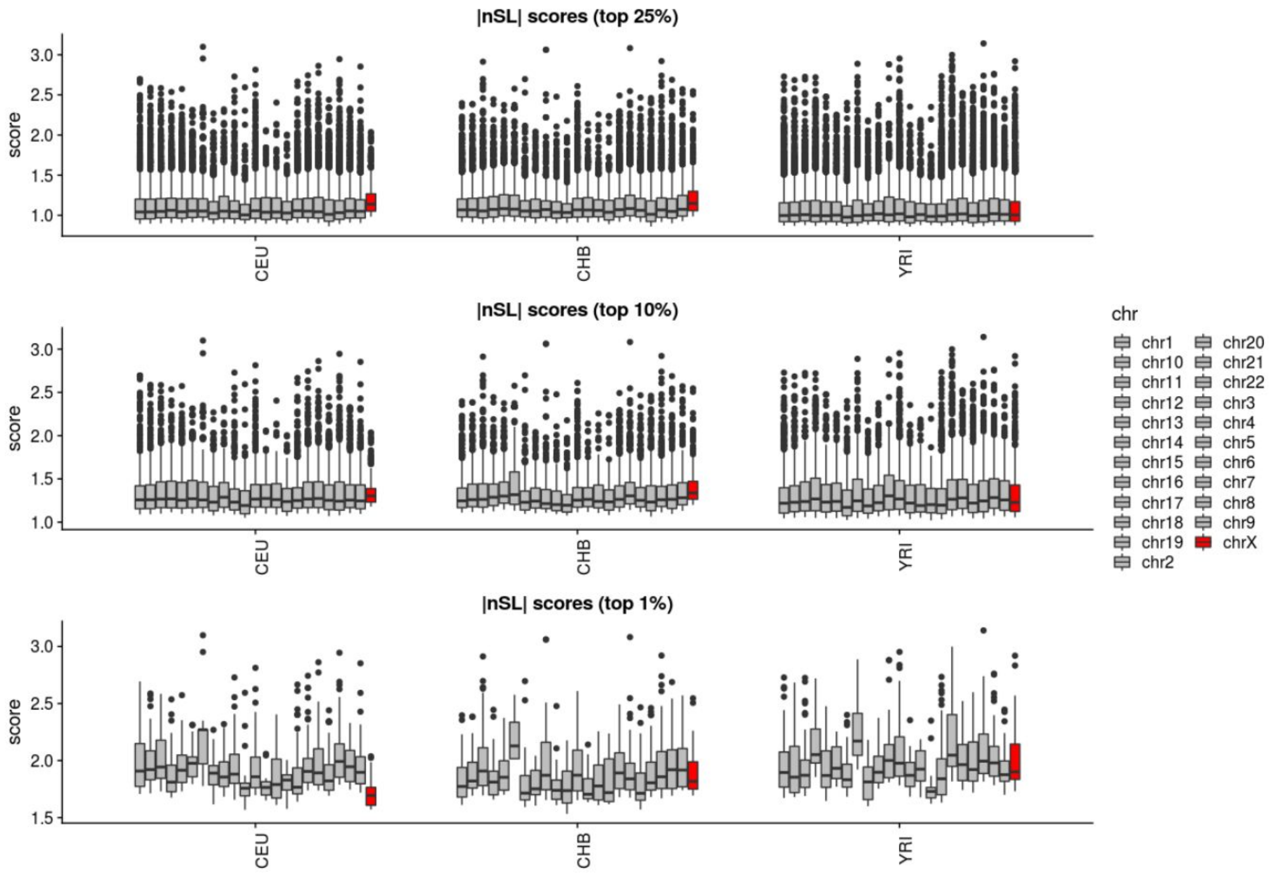


**Supplementary Figure 3.** Comparison between nSL extreme tail distributions of autosomes and X chromosome.

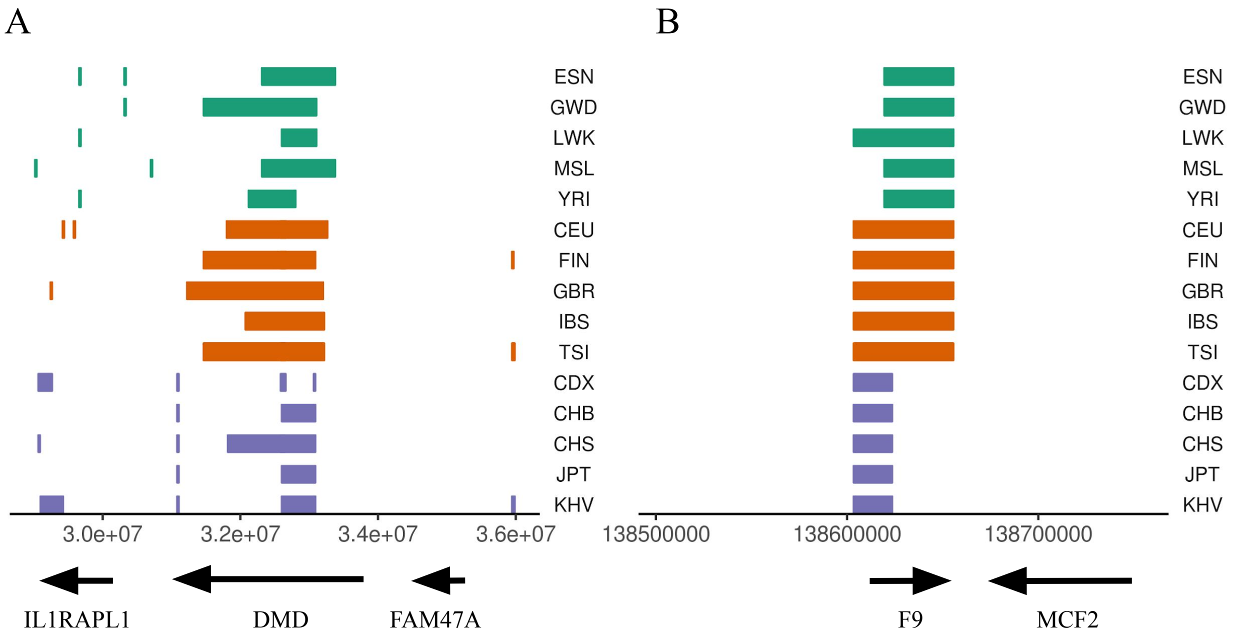


**Supplementary Figure 4.** Sweeping regions under putative positive selection in the three continental groups in the dystrophin gene (*DMD*; A) and coagulation factor 9 (*F9*; B). The reported sweeps are a result of merging the overlapping windows under positive selection in the 99$^{th}$ in all the statistics used (iHS, iHH12 and nSL).

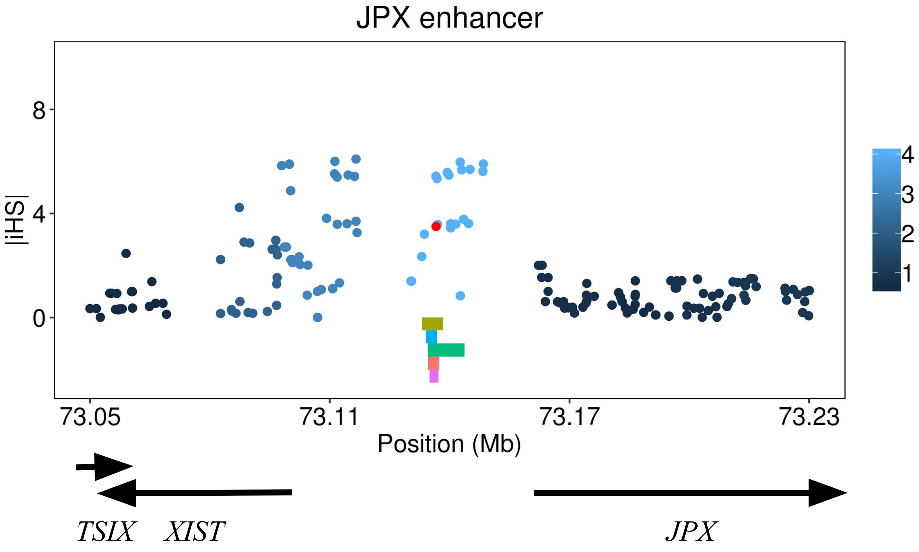


**Supplementary Figure 5.** Manhattan plot showing the putative positive selection signal in African populations reported by iHS in the enhancer located at ~23kb from the *JPX* gene. Marked in red is the SNP rs112977454 reported as eQTL by GTEx. Colour bars at the bottom represent the active enhancers in five different cell lines. Legend shows the window-based score where the SNPs belong to.
