## Supplementary Tables for "Chromosome X-wide analysis of positive selection in human populations: from common and private signals to selection impact on inactivated genes and enhancers-like signatures"

| | | iHS | | nSL | | iHH12 | |
|---|---|---|---|---|---|---|---|
| Group | Population | 99th | 99.9th | 99th | 99.9th | 99th | 99.9th |
| AFR | ESN | 142 | 25 | 185 | 39 | 172 | 51 |
| | GWD | 128 | 32 | 180 | 42 | 166 | 46 |
| | MSL | 153 | 40 | 188 | 39 | 168 | 51 |
| | LWK | 173 | 28 | 190 | 45 | 171 | 59 |
| | YRI | 143 | 31 | 172 | 35 | 181 | 53 |
| EUR | CEU | 61 | 12 | 23 | 0 | 84 | 9 |
| | FIN | 64 | 13 | 29 | 1 | 70 | 15 |
| | GBR | 83 | 13 | 19 | 4 | 88 | 12 |
| | IBS | 58 | 7 | 28 | 8 | 83 | 13 |
| | TSI | 36 | 8 | 21 | 1 | 81 | 12 |
| ASI | CDX | 53 | 9 | 29 | 4 | 57 | 19 |
| | CHB | 53 | 6 | 32 | 3 | 67 | 11 |
| | CHS | 57 | 4 | 28 | 6 | 55 | 12 |
| | JPT | 55 | 7 | 38 | 1 | 54 | 16 |
| | KHV | 56 | 6 | 31 | 4 | 60 | 14 |

**Supplementary Table 1.** Windows under putative positive selection in the extreme simulated $99^{th}$ and $99.9^{th}$ percentiles across the 15 populations under study and the three selection statistics accounting for hard and soft sweeps (iHS, iHH12 and nSL).

| | iHS | | | nSL | | | iHH12 | | |
|---|---|---|---|---|---|---|---|---|---|
| Chr | YRI | CEU | CHB | YRI | CEU | CHB | YRI | CEU | CHB |
| 1 | 180 | 81 | 74 | 229 | 51 | 23 | 277 | 108 | 96 |
| 2 | 191 | 98 | 118 | 325 | 51 | 41 | 273 | 130 | 122 |
| 3 | 228 | 46 | 99 | 205 | 33 | 35 | 224 | 101 | 87 |
| 4 | 193 | 88 | 83 | 275 | 59 | 29 | 207 | 91 | 98 |
| 5 | 146 | 68 | 72 | 248 | 52 | 16 | 205 | 110 | 95 |
| 6 | 119 | 60 | 48 | 230 | 35 | 23 | 239 | 114 | 98 |
| 7 | 123 | 53 | 55 | 190 | 51 | 24 | 180 | 84 | 70 |
| 8 | 120 | 39 | 51 | 202 | 38 | 26 | 178 | 81 | 62 |
| 9 | 69 | 41 | 71 | 161 | 32 | 25 | 132 | 52 | 68 |
| 10 | 123 | 48 | 54 | 156 | 42 | 19 | 145 | 72 | 70 |
| 11 | 120 | 45 | 45 | 169 | 42 | 30 | 165 | 56 | 68 |
| 12 | 86 | 32 | 60 | 196 | 24 | 14 | 170 | 62 | 53 |
| 13 | 72 | 35 | 34 | 130 | 32 | 19 | 124 | 65 | 53 |
| 14 | 79 | 34 | 34 | 117 | 30 | 18 | 109 | 50 | 39 |
| 15 | 66 | 18 | 28 | 85 | 16 | 8 | 66 | 35 | 27 |
| 16 | 61 | 16 | 30 | 95 | 17 | 8 | 87 | 42 | 36 |
| 17 | 50 | 27 | 28 | 63 | 17 | 7 | 71 | 40 | 22 |
| 18 | 65 | 13 | 13 | 95 | 23 | 6 | 92 | 36 | 30 |
| 19 | 31 | 9 | 9 | 73 | 5 | 6 | 54 | 18 | 19 |
| 20 | 41 | 9 | 21 | 67 | 6 | 6 | 65 | 26 | 24 |
| 21 | 31 | 11 | 14 | 45 | 6 | 7 | 42 | 18 | 23 |
| 22 | 27 | 5 | 6 | 33 | 7 | 6 | 34 | 11 | 12 |

**Supplementary Table 2.** Windows under putative positive selection in the extreme simulated 99$^{th}$ percentiles in the human autosomes of the three populations of reference (YRI, CEU and CHB) across the three selection statistics accounting for hard and soft sweeps (iHS, iHH12 and nSL).

| Population | Test | GO term | Description | FDR |
|---|---|---|---|---|
| ESN | iHS,nSL | MF:GO:0004065 | sulfuric ester hydrolase activity | 0.00074 |
| | iHS,nSL | CC:GO:0004065 | endoplasmic reticulum lumen | 0.046 |
| | iHH12 | CC:GO:0045211 | postsynaptic membrane | 0.017 |
| | iHH12 | CC:GO:1902495 | transmembrane transporter complex | 0.017 |
| | iHH12 | CC:GO:0030425 | dendrite | 0.0475 |
| | nSL | MF:GO:0042043 | neurexin family protein binding | 0.0085 |
| GWD | iHS | MF:GO:0008484 | sulfuric ester hydrolase activity | 0.00000117 |
| | iHS | CC:GO:0005587 | collagen type IV trimer | 0.032 |
| | iHS | CC:GO:0031968 | organelle outer membrane | 0.023 |
| | iHS | CC:GO:0044432 | endoplasmic reticulum part | 0.0093 |
| | iHH12 | CC:GO:0005884 | actin filament | 0.039 |
| | iHH12 | CC:GO:1902495 | transmembrane transporter complex | 0.021 |
| | iHH12 | CC:GO:0044463 | cell projection part | 0.0217 |
| | nSL | MF:GO:0004065 | arylsulfatase activity | 0.021 |
| | nSL | MF:GO:0042043 | neurexin family protein binding | 0.0059 |
| MSL | iHS | MF:GO:0008484 | sulfuric ester hydrolase activity | 0.0000043 |
| | iHH12,nSL | MF:GO:0004065 | arylsulfatase activity | 0.0197 |
| | iHH12 | CC:GO:0097060 | synaptic membrane | 0.0163 |
| | iHH12 | CC:GO:1902495 | transmembrane transporter complex | 0.0381 |
| | iHH12 | CC:GO:0044463 | cell projection part | 0.0381 |
| LWK | iHS | MF:GO:0008484 | sulfuric ester hydrolase activity | 0.000653 |
| | iHS | CC:GO:0005788 | endoplasmic reticulum lumen | 0.00617 |
| | iHH12 | CC:GO:0045211 | postsynaptic membrane | 0.032 |
| | iHH12 | CC:GO:1902495 | transmembrane transporter complex | 0.039 |
| | iHH12 | CC:GO:0030425 | dendrite | 0.039 |
| | nSL | MF:GO:0004065 | arylsulfatase activity | 0.0169 |
| YRI | iHS | MF:GO:0008484 | sulfuric ester hydrolase activity | 0.000610 |
| | iHS | CC:GO:0005788 | endoplasmic reticulum lumen | 0.0116 |
| | iHS | CC:GO:0043005 | neuron projection | 0.00086 |
| | iHS | CC:GO:0045202 | synapse | 0.0026 |
| | nSL | MF:GO:0042043 | neurexin family protein binding | 0.016 |

**Supplementary Table 3A.** Significant GO terms of the top 100 genes across all the Sub-saharan African populations in the three selection tests used in the analysis. We consider FDR < 0.05 as significant. In the table, we present the population ID, the tests where the term is reported as significant, the GO term ID, the term description and the corrected FDR value.

| Population | Test | GO term | Description | FDR |
|---|---|---|---|---|
| CEU | iHS,nSL | BP:GO:0097105 | presynaptic membrane assembly | 0.025 |
| | iHS,nSL | MF:GO:0042043 | neurexin family protein binding | 0.0122 |
| | iHS,nSL | CC:GO:0098985 | asymmetric, glutamatergic, excitatory synapse | 0.0179 |
| | iHS,iHH12,nSL | CC:GO:0097060 | synaptic membrane | 0.0053 |
| | iHS | CC:GO:0030425 | dendrite | 0.0182 |
| | iHH12,nSL | CC:GO:0044456 | synapse part | 0.0011 |
| | iHH12 | CC:GO:0044463 | cell projection part | 0.013 |
| | nSL | BP:GO:0042391 | regulation of membrane potential | 0.019 |
| FIN | iHS | CC:GO:0000775 | chromosome, centromeric region | 0.034 |
| | iHS,nSL | CC:GO:0030425 | dendrite | 0.003 |
| | iHS | CC:GO:0043005 | neuron projection | 0.003 |
| | iHH12 | CC:GO:0097060 | synaptic membrane | 0.0071 |
| | iHH12 | CC:GO:0044463 | cell projection part | 0.0085 |
| | nSL | MF:GO:0042043 | neurexin family protein binding | 0.0109 |
| | nSL | CC:GO:0098985 | asymmetric, glutamatergic, excitatory synapse | 0.0232 |
| | nSL | CC:GO:0036020 | endolysosome membrane | 0.0437 |
| | nSL | BP:GO:0097105 | presynaptic membrane assembly | 0.0211 |
| GBR | iHS | CC:GO:0030425 | dendrite | 0.0016 |
| | iHS | CC:GO:0097060 | synaptic membrane | 0.0183 |
| | iHS | CC:GO:0043005 | neuron projection | 0.001 |
| | iHH12 | CC:GO:0043296 | apical junction complex | 0.01 |
| | iHH12 | CC:GO:0030426 | growth cone | 0.0155 |
| | iHH12 | CC:GO:1902495 | transmembrane transporter complex | 0.033 |
| | nSL | BP:GO:0097105 | presynaptic membrane assembly | 0.049 |
| | nSL | CC:GO:0097458 | neuron part | 0.0041 |
| | nSL | MF:GO:0042043 | neurexin family protein binding | 0.019 |
| IBS | iHS | CC:GO:0097060 | synaptic membrane | 0.03 |
| | iHH12 | CC:GO:0150034 | distal axon | 0.012 |
| | iHH12 | CC:GO:1902495 | transmembrane transporter complex | 0.024 |
| | iHH12 | CC:GO:0098794 | postsynapse | 0.0006 |
| TSI | iHS | CC:GO:0030425 | dendrite | 0.0007 |
| | iHS | CC:GO:0044456 | synapse part | 0.002 |
| | iHS | CC:GO:0043005 | neuron projection | 0.0007 |
| | iHH12 | CC:GO:1902495 | transmembrane transporter complex | 0.0177 |
| | iHH12 | CC:GO:0030425 | dendrite | 0.0047 |
| | iHH12 | CC:GO:0045211 | postsynaptic membrane | 0.002 |
| | nSL | CC:GO:0097458 | neuron part | 0.016 |

**Supplementary Table 3B.** Significant GO terms of the top 100 genes across all the European populations in the three selection tests used in the analysis. We consider FDR < 0.05 as significant. In the table, we present the population ID, the tests where the term is reported as significant, the GO term ID, the term description and the corrected FDR value.

| Population | Test | GO term | Description | FDR |
|---|---|---|---|---|
| CDX | iHS,iHH12,nSL | BP:GO:0097105 | presynaptic membrane assembly | 0.03 |
| | iHS,iHH12,nSL | CC:GO:0005587 | collagen type IV trimer | 0.029 |
| | iHS,iHH12,nSL | CC:GO:0098985 | asymmetric, glutamatergic, excitatory synapse | 0.029 |
| | iHS | CC:GO:0098794 | postsynapse | 0.026 |
| | iHH12,nSL | CC:GO:0045202 | synapse | 0.025 |
| CHB | iHH12,nSL | BP:GO:0097105 | presynaptic membrane assembly | 0.032 |
| | iHH12 | CC:GO:0098982 | GABA-ergic synapse | 0.034 |
| | iHH12 | CC:GO:0005794 | Golgi apparatus | 0.034 |
| CHS | iHS,iHH12,nSL | BP:GO:0097105 | presynaptic membrane assembly | 0.0497 |
| | iHS,nSL | CC:GO:0098794 | postsynapse | 0.0296 |
| | iHS | CC:GO:0106018 | phosphatidylinositol-3,5-bisphosphate phosphatase activity | 0.0296 |
| | iHH12 | CC:GO:0098985 | asymmetric, glutamatergic, excitatory synapse | 0.0197 |
| | iHH12 | CC:GO:0030425 | dendrite | 0.028 |
| | iHH12 | CC:GO:0044456 | synapse part | 0.011 |
| JPT | iHS,iHH12 | CC:GO:0005587 | collagen type IV trimer | 0.049 |
| | iHS | CC:GO:0099086 | synaptonemal structure | 0.049 |
| | iHS,iHH12 | CC:GO:0005794 | Golgi apparatus | 0.049 |
| | iHH12,nSL | BP:GO:0097105 | presynaptic membrane assembly | 0.0342 |
| | iHH12 | CC:GO:0098985 | asymmetric, glutamatergic, excitatory synapse | 0.023 |
| | iHH12 | CC:GO:0098794 | postsynapse | 0.021 |
| | nSL | MF:GO:0042043 | neurexin family protein binding | 0.0188 |
| KHV | iHS,iHH12,nSL | BP:GO:0097105 | presynaptic membrane assembly | 0.036 |
| | iHS,nSL | CC:GO:0005587 | collagen type IV trimer | 0.036 |
| | iHS,nSL | CC:GO:0098985 | asymmetric, glutamatergic, excitatory synapse | 0.036 |
| | iHS | CC:GO:0045211 | postsynaptic membrane | 0.036 |
| | nSL | CC:GO:0098794 | postsynapse | 0.029 |

**Supplementary Table 3C.** Significant GO terms of the top 100 genes across all the Asian populations in the three selection tests used in the analysis. We consider FDR < 0.05 as significant. In the table, we present the population ID, the tests where the term is reported as significant, the GO term ID, the term description and the corrected FDR value.

| | | iHS | | | iHH12 | | | nSL | | |
|---|---|---|---|---|---|---|---|---|---|---|
| | | Selected | Not Sel. | Total | Selected | Not Sel. | Total | Selected | Not Sel. | Total |
| **95th** | **Escape** | 31 | 28 | 59 | 17 | 42 | 59 | 18 | 41 | 59 |
| | **Inactive** | 133 | 248 | 381 | 68 | 313 | 381 | 136 | 245 | 381 |
| | **Total** | 164 | 276 | 440 | 85 | 355 | 440 | 154 | 286 | 440 |
| **99th** | **Escape** | 18 | 41 | 59 | 10 | 49 | 59 | 9 | 50 | 59 |
| | **Inactive** | 44 | 337 | 381 | 34 | 347 | 381 | 50 | 331 | 381 |
| | **Total** | 62 | 378 | 440 | 44 | 396 | 440 | 59 | 381 | 440 |
| **99.9th** | **Escape** | 8 | 51 | 59 | 3 | 56 | 59 | 4 | 55 | 59 |
| | **Inactive** | 14 | 367 | 381 | 10 | 371 | 381 | 12 | 369 | 381 |
| | **Total** | 22 | 418 | 440 | 13 | 427 | 440 | 16 | 424 | 440 |

**Supplementary Table 4A.** Contingency tables of escape genes under selection reported by the three selection statistics across three extreme percentiles (95th, 99th and 99.9th). Two categories were used: escape/inactive and selected/non-selected.

| | iHS | | | iHH12 | | | nSL | | |
|---|---|---|---|---|---|---|---|---|---|
| | Fisher's p | O.R | C.I.(0.95) | Fisher's p | O.R. | C.I.(0.95) | Fisher's p | O.R. | C.I.(0.95) |
| **95th** | 0.01 | 2.06 | 1.14-3.73 | 0.05 | 1.86 | 0.93-3.57 | 0.46 | 0.79 | 0.41-1.47 |
| **99th** | 0.0003 | 3.35 | 1.66-6.59 | 0.06 | 2.07 | 0.86-4.64 | 0.68 | 1.19 | 0.48-2.64 |
| **99.9th** | 0.004 | 4.09 | 1.41-11.07 | 0.40 | 1.98 | 0.34-8.02 | 0.24 | 2.23 | 0.50-7.70 |

**Supplementary Table 4B.** Fisher's tests applied to the contingency tables. iHS reports significant p-values across the three extreme percentiles with increasing odds ratios (OR). iHH12 and nSL do not show significant enrichment in escape genes, however the odds are in line with those in iHS in five out of the six comparisons, suggesting the presence of selection but with lack of significance probably due to a sample effect.

| | AFR | EUR | ASI |
|---|---|---|---|
| **95th** | AP1S2, ARSD, ARSE, ARSF, ARSH, CDK16, FAM9C, HS6ST2, HTR2C, JPX, KDM6A, MAGEC3, MAOA, MED14, MSL3, MXRA5, NR0B1, OFD1, PCDH19, PNPLA4, PRKX, STS, TMEM27, UBA1, ZCCHC16, ZRSR2 | ARSF, GYG2, HS6ST2, HTR2C, MAGEC3, STS, TAF7L, TMEM27, USP9X, ZCCHC16, ZFX | FUNDC1, KDM6A, STS, USP9X |
| **99th** | ARSE, ARSF, ARSH, CDK16, FAM9C, HS6ST2, HTR2C, KDM6A, MAGEC3, MAGEC3, MED14, MXRA5, OFD1, STS, TMEM27, UBA1, | HS6ST2, STS, USP9X, ZCCHC16 | FUNDC1,KDM6A, STS |
| **99th** | ARSF, ARSH, CDK16, HTR2C, KDM6A, STS, UBA1 | ZCCHC16 | - |

**Supplementary Table 4C.** Escape genes reported by iHS as being under positive selection in each continental group across the extreme percentiles.

| Group | Population | iHS | | | | iHH12 | | | | nSL | | | |
|---|---|---|---|---|---|---|---|---|---|---|---|---|---|
| | | Int | I | E | D | Int | I | E | D | Int | I | E | D |
| AFR | ESN | 18 | 5 | 0 | 0 | 30 | 16 | 1 | 0 | 17 | 6 | 0 | 0 |
| | GWD | 18 | 11 | 1 | 0 | 34 | 30 | 2 | 0 | 19 | 9 | 0 | 1 |
| | MSL | 18 | 19 | 0 | 0 | 39 | 34 | 3 | 0 | 17 | 9 | 0 | 0 |
| | LWK | 32 | 15 | 0 | 0 | 56 | 35 | 2 | 0 | 18 | 10 | 0 | 0 |
| | YRI | 14 | 10 | 1 | 0 | 38 | 18 | 1 | 0 | 18 | 7 | 0 | 0 |
| EUR | CEU | 3 | 1 | 0 | 0 | 12 | 3 | 0 | 0 | 0 | 0 | 0 | 0 |
| | FIN | 1 | 4 | 1 | 0 | 11 | 6 | 0 | 0 | 0 | 0 | 0 | 0 |
| | GBR | 3 | 1 | 0 | 0 | 19 | 2 | 0 | 0 | 4 | 1 | 0 | 0 |
| | IBS | 2 | 3 | 0 | 0 | 9 | 7 | 0 | 0 | 5 | 1 | 0 | 0 |
| | TSI | 3 | 5 | 0 | 0 | 6 | 7 | 0 | 0 | 0 | 1 | 0 | 0 |
| ASI | CDX | 5 | 1 | 0 | 0 | 17 | 4 | 0 | 0 | 1 | 1 | 0 | 0 |
| | CHB | 1 | 0 | 0 | 0 | 15 | 0 | 0 | 0 | 4 | 0 | 0 | 0 |
| | CHS | 4 | 0 | 0 | 0 | 16 | 0 | 0 | 0 | 4 | 0 | 0 | 0 |
| | JPT | 1 | 0 | 0 | 0 | 16 | 1 | 0 | 0 | 0 | 0 | 0 | 0 |
| | KHV | 3 | 2 | 0 | 0 | 17 | 3 | 0 | 0 | 0 | 0 | 0 | 0 |

**Supplementary Table 5.** SNPs with a selection score within the 1% extreme and with CADD score ≥ 10 in the 99.9th percentile across all populations (Intergenic (Int), Intronic (I), Exonic (E), Downstream (D)

| Test | Region | n SNPs | RegulomeDB scores | | | | | ENCODE elements | OR |
|---|---|---|---|---|---|---|---|---|---|
| | | | 2 | 3 | 4 | 5 | 6 | | |
| iHS | Intergenic | 31 (0.1) | - | - | 1 | 1 | 9 | 35% | 1.08 |
| | Genic | 286 (0.9) | - | - | 6 | 16 | 74 | 33% | |
| iHH12 | Intergenic | 35 (0.14) | - | - | 1 | 6 | 11 | 51% | 1.32 |
| | Genic | 212 (0.85) | 2 | 2 | 8 | 25 | 57 | 44% | |
| nSL | Intergenic | 32 (0.12) | - | - | - | 3 | 10 | 40% | 1.23 |
| | Genic | 230 (0.88) | 1 | 4 | 4 | 9 | 64 | 35% | |

**Supplementary Table 6A.** RegulomeDB annotation of the 99th percentile genic windows with intergenic overlap and the odds ratio (OR) between genic and intergenic SNPs within functional elements. All populations are considered.

| Test | Region | n SNPs | RegulomeDB scores | | | | | ENCODE elements | OR |
|---|---|---|---|---|---|---|---|---|---|
| | | | 2 | 3 | 4 | 5 | 6 | | |
| iHS | Intergenic | 4 (0.05) | - | - | - | - | 1 | 25% | 0.77 |
| | Genic | 83 (0.95) | - | - | 3 | 3 | 19 | 30% | |
| iHH12 | Intergenic | 8 (0.08) | - | - | - | 2 | 5 | 87% | 8.24 |
| | Genic | 98 (0.92) | 1 | 1 | 2 | 15 | 26 | 45% | |
| nSL | Intergenic | 10 (0.16) | - | - | - | 1 | 5 | 60% | 2.11 |
| | Genic | 41 (0.84) | 1 | - | 2 | 1 | 13 | 41% | |

**Supplementary Table 6B.** RegulomeDB annotation of the 99th percentile genic windows with intergenic overlap when considering extreme scoring SNPs (per-SNP 1% extreme tail). iHH12 and nSL show a significant OR increment in comparison with iHS

| Test | Group | 99th | | | | 99.9th | | | |
|---|---|---|---|---|---|---|---|---|---|
| | | Non-ovlp | Ovlp. | % | OR | Non-ovlp | Ovlp. | % | OR |
| iHS | AFR | 194 | 3 | 1.52 | 0.46 | 47 | 3 | 6.00 | 1.93 |
| | EUR | 116 | 1 | 0.86 | 0.26 | 23 | 0 | 0 | 0 |
| | ASI | 111 | 7 | 5.93 | 1.93 | 17 | 0 | 0 | 0 |
| iHH12 | AFR | 128 | 17 | 11.72 | 4.31 | 43 | 3 | 6.52 | 2.11 |
| | EUR | 77 | 4 | 4.94 | 1.57 | 17 | 2 | 10.52 | 3.57 |
| | ASI | 79 | 8 | 9.20 | 3.14 | 22 | 1 | 4.35 | 1.37 |
| nSL | AFR | 283 | 4 | 1.39 | 0.41 | 68 | 2 | 2.86 | 0.88 |
| | EUR | 52 | 1 | 1.89 | 0.58 | 8 | 1 | 11.11 | 3.77 |
| | ASI | 77 | 2 | 2.53 | 0.78 | 8 | 0 | 0 | 0 |

**Supplementary Table 7.** Overlapping and non-overlapping intergenic windows under putative positive selection on enhancer regions reported by HACER in any cell line (see Methods) across the three continental groups. Odds ratio (OR) of intergenic and overlapping windows shows a significant increment mainly in iHH12 across all populations.

| | | Observed selection on target gene | | |
|---|---|---|---|---|
| | | Y | N | Total |
| Observed selection on enhancer | Y | 218 | 350 | 568 |
| | N | 167 | 395 | 562 |
| | Total | 385 | 745 | 1130 |

| | | Expected selection on target gene | | |
|---|---|---|---|---|
| | | Y | N | Total |
| Expected selection on enhancer | Y | 193 | 375 | 568 |
| | N | 192 | 370 | 562 |
| | Total | 385 | 745 | 1130 |

**Supplementary Table 8.** Contingency tables of both observed and expected pairs of enhancer/target-gene in the following categories: Selected enhancer and selected gene (YY), Selected enhancer and non-selected gene (YN), Non-selected enhancer and selected gene (NY), Non-selected enhancer and non-selected gene (NN). A Chi square test is applied to study the dependency of both variables (Chi sqr value = 9.44; p-value = 0.0021).

| Chr | Start | End | Length | Test | Population | Closest gene |
|---|---|---|---|---|---|---|
| X | 40238664 | 40241510 | 2846 | iHH12 | CEU,GBR,IBS | ATP6AP2 |
| X | 45179990 | 45196717 | 16727 | iHH12 | FIN,TSI | KDM6A(*) |
| X | 53740262 | 53744843 | 4581 | nSL | GWD | HUWE1(**) |
| X | 73135561 | 73145161 | 9600 | iHS | ESN,GWD,MSL,LWK,YRI | JPX |
| X | 109017803 | 109018393 | 590 | iHS,nSL | YRI | ACSL4(**) |
| X | 123351438 | 123353650 | 2212 | iHH12 | GWD,MSL,YRI | SH2D1A(**) |

**Supplementary Table 9.** Top enhancer regions under putative positive selection (99.9$^{th}$ percentile). The genes marked as "**" are found under selection in sequence in the 99.9$^{th}$ percentile and in the same continental group, the genes marked as "*" are found under selection as well but in a different continental group.
